# Behavioral rhythms are bad predictors of general organismal rhythmicity

**DOI:** 10.1101/2022.07.16.500291

**Authors:** N. Sören Häfker, Laurenz Holcik, Karim Vadiwala, Isabel Beets, Audrey M. Mat, Alexander W. Stockinger, Stefan Hammer, Liliane Schoofs, Florian Raible, Kristin Tessmar-Raible

## Abstract

The circadian clock controls behavior and metabolism in various organisms. However, the exact timing and strength of rhythmic phenotypes can vary significantly between individuals of the same species. This is highly relevant for the rhythmically complex marine environments where organismal rhythmic diversity likely permits the occupation of different microenvironments. When investigating circadian locomotor behavior of *Platynereis dumerilii,* a model system for marine molecular chronobiology, we found strain-specific, high variability between individual worms. The individual patterns were reproducibly maintained for several weeks independent of basic culture conditions, such as population density or feeding. A diel head transcriptome comparison of behaviorally rhythmic versus arrhythmic wildtype worms showed that 24h cycling of core circadian clock transcripts is identical between both behavioral phenotypes. While behaviorally arrhythmic worms showed a similar total number of cycling transcripts compared to their behaviorally rhythmic counterparts, the annotation categories of their transcripts, however, differed substantially. Consistent with their locomotor phenotype, behaviorally rhythmic worms exhibit an enrichment of cycling transcripts related to neuronal/behavioral processes. In contrast, behaviorally arrhythmic worms showed significantly increased diel cycling for metabolism- and physiology-related transcripts. Phenotype-specific cycling of distinct matrix metalloproteinase transcripts, encoding extracellular enzymes that modulate synaptic circuit function and neuropeptide signaling, like pigment dispersing factor (PDF), prompted us to functionally investigate *Platynereis pdf*. Differing from its role in *Drosophila,* loss of *pdf* impacts on overall activity levels, but shows only indirect effects on rhythmicity. Our results show that individuals arrhythmic in a given process can show increased rhythmicity in others. Across the *Platynereis* population, variations of this exist as a reproducible continuum. We hypothesize that such diel rhythm breadth is an important biodiversity resource enabling the species to quickly adapt to heterogeneous marine environments and potentially also to the effects of climate change, which is however endangered with shrinking population sizes and hence diversity.

## INTRODUCTION

Life on earth has been evolving in the presence of numerous environmental cycles. This is particularly prominent in the marine environment [1, 2], where rhythms even exist in the deep sea [3]. In the oceans, biological rhythms can be the major factors shaping ecosystem structure and productivity, and they significantly contribute to biogeochemical cycling [4–6], but compared to the terrestrial environment our understanding of the factors and mechanisms driving marine rhythmicity is still severely limited [2]. This is particularly concerning in the context of climate change as it is largely unknown how marine timing systems and associated species fitness on the individual and population level will be affected by changing environmental conditions [2, 7].

Rhythmic adaptations on the level of behavior, physiology and gene expression are central in shaping an animal‟s interaction with the abiotic and biotic environment. Many of these rhythms are under the control of endogenous timing mechanisms (clocks) [7, 8]. By far best studied is the circadian clock that enables organisms to synchronize (entrain) to, and to anticipate the 24 hour day/night cycle [9], thereby significantly contributing to fitness [10, 11]. Like all biological processes, ∼24hr rhythmicity varies between individuals, as prominently illustrated by human chronotypes [12]. Studies in terrestrial organisms, like drosophilid flies, identified differences in the expression of circadian clock genes and the neuropeptide pigment-dispersing factor (PDF) to be responsible for species- and population-specific patterns of diel activity [13–18]. Similarly, differences in clock gene expression rhythms were linked to caste-specific activity phenotypes in ants [19]. Comparisons of fly strains and species from different latitudes further indicate that behavioral rhythmicity depends on the localization of circadian core clock genes expression in the brain [16, 20] and clock gene alleles [21, 22]. The marine midge *Clunio* shows habitat- and strain-specific temporal niche adaptations of circadian emergence timing that are linked to splice variants in a *calcium-calmodulin-dependent kinase*. These splice variants impact on core clock protein phosphorylation thereby affecting circadian period length [23]. Furthermore, behavioral phenotypes including arrhythmicity can also depend on the circadian clock output pathways [17]. It is often assumed that the rhythm power of a particular read-out, e.g. locomotor activity, is representative for the rhythmicity of the entire organism, but integrative studies that test this assumption are rare.

Behavioral variability can arise seemingly at random, but can also be consistent over time, then referred to as „personality‟, „behavioral syndrome‟ or „coping style‟[24, 25]. While often such behavioral „personalities‟ are intuitively associated to vertebrates, they have in fact been documented in a variety of organisms from invertebrates to mammals [24–26]. Due to the relatively short life time of *Drosophila melanogaster*, individual variance of ∼24hr rhythms can only be explored over a limited time and individuals cannot be re-tested weeks later. However, many invertebrates live much longer, and especially in the context of understanding the adaptation potential of circadian clocks in populations, comprehending the extent and source of their variability is critical. The understanding of how differences in circadian behavioral rhythms across individuals and populations interlink with their molecular features are still highly limited. This applies especially to marine organisms, despite the fundamental importance of biological rhythms and clocks for marine ecosystems [2].

The marine polychaete *Platynereis dumerilii* is a molecularly slowly evolving annelid that has emerged as a functional model for marine chronobiology with a life cycle representative for various marine invertebrates [27–30]. After a planktonic larvae phase during which animals get dispersed by currents, the worm switches to a benthic lifestyle that ends with a metamorphic maturation into a free-swimming spawning form [27]. It inhabits temperate to tropical shores worldwide, typically at depths of 0-10 m [31, 32]. The experimental culture used here derives from the Mediterranean, where the worm is typically part of the ecologically important *Posedonia* sea grass meadows [30].

Here, we explore ∼24hr behavioral diversity of *Platynereis* strains and individuals, its transcriptome-wide consequences and the role of PDF, an important neuropeptide for behavioral rhythmicity in *Drosophila melanogaster* [13, 14]. We asked, if the behavioral circadian variability in *Platynereis* occurs at random or represents a reproducible individual trait, and how this might be reflected on the molecular level. Worms displayed strain-specific characteristics as well as strong and reproducible inter-individual variability in diel behavior, even among co-raised siblings. A diel transcriptome comparison of behaviorally rhythmic versus arrhythmic wildtype worms showed that these phenotypic differences match with differences in diel cyclic transcripts. This was especially evident for transcripts involved in neuronal signaling versus metabolic processes. In contrast, cycling of core circadian clock genes was identical between the different behavioral groups. The transcriptomic analyses further pointed towards a potential phenotype-specific role of *matrix metalloproteinases* (*mmps*) in the establishment of individual behavioral patterns. Given that in *Drosophila,* MMP1 mediates structural plasticity of PDF+ clock neuron circuits, and thereby diel behavior, in a PDF-dependent manner [33, 34], we tested the effects of *pdf* knock-outs in *Platynereis.* The phenotype was remarkably different from *Drosophila,* as there was no direct effect on circadian period or rhythms power of locomotor activity after several outcrosses. Instead, *pdf* mutants showed overall more activity, which occasionally had indirect effects on rhythm power under light/dark-cycles. This suggests that in *Platynereis* the level of behavioral versus metabolic rhythm power is controlled by genes downstream of the core circadian clock (e.g. *mmps*, gene of neuronal functions). Our work also provides new perspectives on the evolution of PDF. While PDF’s role in general behavioral control is conserved across major invertebrate groups, its role in circadian control has to be questioned outside insects.

Considering the complexity of the cyclic environment in marine habitats [1, 2], unraveling the molecular processes shaping phenotypic rhythms and their diversity will be important to understand how marine species adapt to these conditions, and how individuals and populations cope with the ongoing shifts in habitat conditions due to climate change.

## RESULTS

### *Platynereis* circadian behavioral rhythmicity differs between strains, but is individually consistent and reproducible

In our previous studies we repeatedly observed a high diversity in circadian behaviors of immature and premature *Platynereis dumerilii* bristle worms [28,35,36]. This prompted us to systematically investigate the worms‟ diversity in circadian locomotor rhythmicity patterns. Please note that below we use the term „rhythmicity/arrhythmicity‟ to specifically refer to behavior, while we use the term „cycling‟ in the context of gene expression.

We chose three worm strains (VIO & PIN: collected from the Bay of Naples, Italy more than 50 years ago and continuously incrossed over the past 14 years; NAP: newly collected from the Bay of Naples in 2009 and maintained as a non-incrossed, closed culture) [30, 37]. We repeatedly recorded locomotor behavior of immature/premature sibling worms for 4 days of 16h:8h light/dark-cycle (LD) followed by 4 days of constant darkness (DD) (Fig 1A, S1 Fig). All strains were generally nocturnal and showed little overall inter-strain differences in morning/afternoon/night activity (Fig 1B). However, when we analyzed rhythmicity (20h-28h period range) under LD and DD in more detail, we noticed that strain-specific rhythmicity patterns and rhythm power varied significantly between strains (Fig 1A,C,E-G). Aside the inter-strain differences, all strains displayed pronounced inter-individual variability in rhythm power that is best described as a behavioral continuum reaching from highly rhythmic to fully arrhythmic worms (Fig 1C,D, S2 Fig).

**Fig 1:**
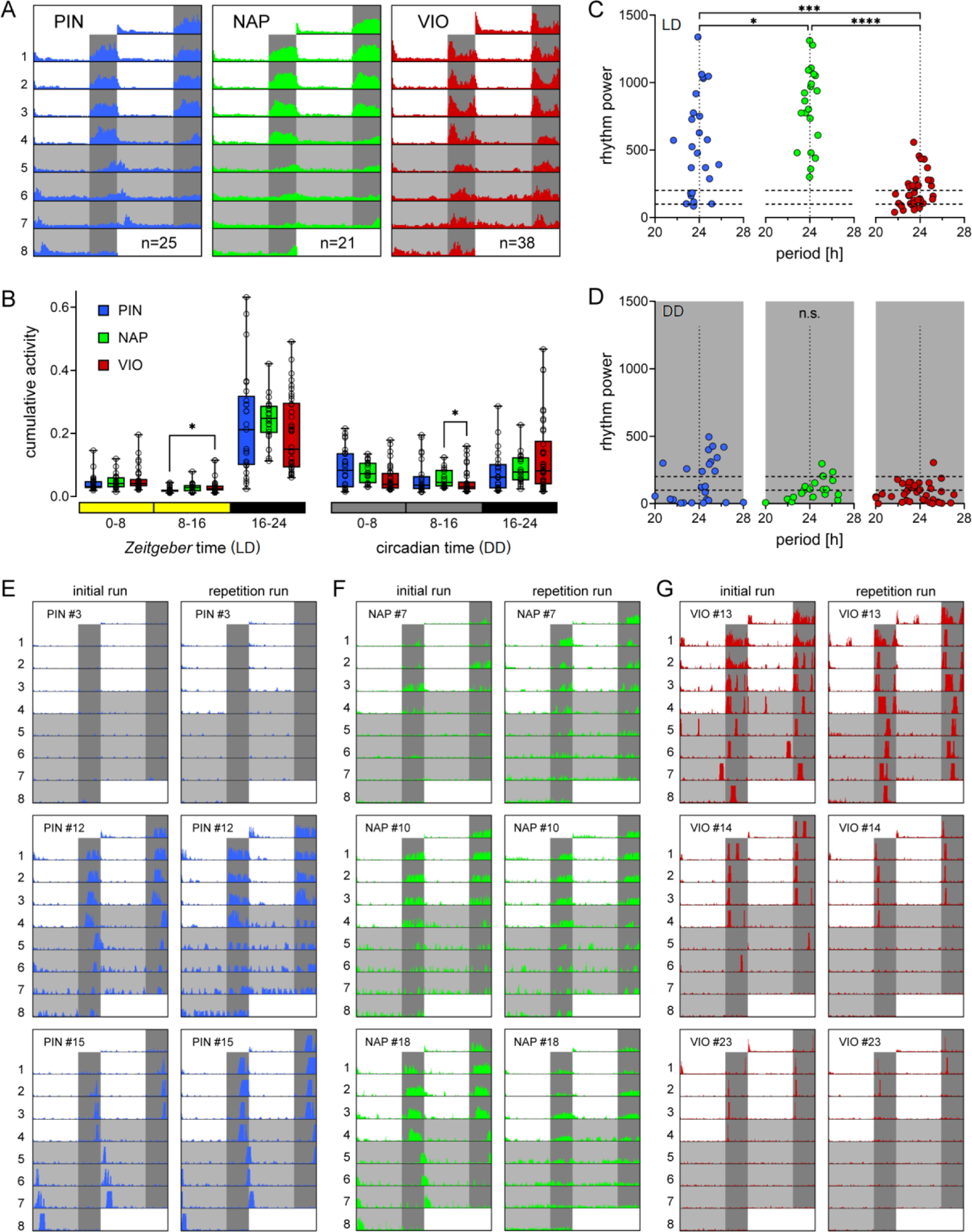
Inter-strain and inter-individual variability in circadian behavioral rhythmicity. The circadian locomotor activities of the three *Platynereis* strains PIN (blue), NAP (green) and VIO (red) are compared. (A) Double-plotted actograms of mean locomotor activity over 4 days of a 16h:8h light/dark-cycle (LD) followed by 4 days of constant darkness (DD). Individual worm actograms are provided in S2 Fig. (B) Cumulative activity across early day (0-8), late day (8-16) and night (16-24) in LD and DD. Box: median with 25%/75% percentiles, whiskers: min/max. (C,D) Period/power of locomotor rhythms in the 20h-28h range determined by Lomb-Scargle periodogram. Dashed horizontal lines: thresholds for weak rhythm power (<200) and arrhythmicity (<100) based on visual actogram inspection. (E-G) Exemplary actograms of PIN, NAP and VIO individuals investigated in two consecutive runs (initial/repeated). #: individual worm identifier. Statistical differences were determined via Kruskal-Wallis ANOVA on Ranks (panels B,D) or Welch‟s ANOVA (panel C). Significance levels: \**p*<0.05, \*\**p*<0.01, \*\*\**p*<0.001, \*\*\*\**p*<0.0001. Each circle: individual worm. For additional initial/repeated actograms see S3 Fig.

To closer investigate the observed individual diversity, we performed repetition runs on a total of 30 worms (PIN n= 12, NAP n=5, VIO n=13) 1-2 months after the initial recordings. Repetitions were limited due to maturation of the other individuals. Actograms of the first and second recordings were very reproducible showing the same behavioral characteristics in both runs. The highest diversity of behavioral patterns was observed in PIN worms, which, however, also exhibited the highest consistency between initial and repeated runs (Fig 1E-G, S3 Fig). Of note, after the initial run we kept all worms individually for identification, which are very different conditions from the “group maintenance” that the worms experienced before the initial run. This shows that the diverse circadian activity patterns in *Platynereis* are highly reproducible in individuals over time, irrespective of basic culture conditions.

### Core circadian clock gene transcripts cycle identically in rhythmic and arrhythmic worms

In order to obtain insight into the molecular characteristics associated with the individually different rhythmic behavior, we investigated the diel transcriptome of behaviorally rhythmic and arrhythmic wildtype worms. We focused on worms of the PIN strain, as their circadian activity patterns showed the strongest within-individual reproducibility (Fig 1E, FigS3A).

Behavior of wildtype worms was recorded for 3 LD days and 3 DD days, and worms were characterized as either rhythmic (clear 24h rhythmicity), arrhythmic (no rhythmicity) or intermediate (everything in between) based on visual actogram inspection and ActogramJ rhythm analysis (Fig 2A, S4 Fig). Heads of worms identified as behaviorally rhythmic or arrhythmic were collected under LD in 4h intervals starting at ZT0 (lights on). Transcriptome sequencing on a NextSeq550 System (Illumina, USA) resulted in a total of 445,553,522 raw reads (Fig 2B). Trimming/filtering caused a 0.48% read loss and trimmed reads were mapped against an independently generated transcriptome (see methods). The mapping results were filtered to exclude transcripts that were not/hardly expressed and/or <500bp, resulting in 48,605 distinct expressed transcripts (S1 Tab), highly consistent with previous differential expression by sequencing (DEseq) analyses [38].

**Fig 2:**
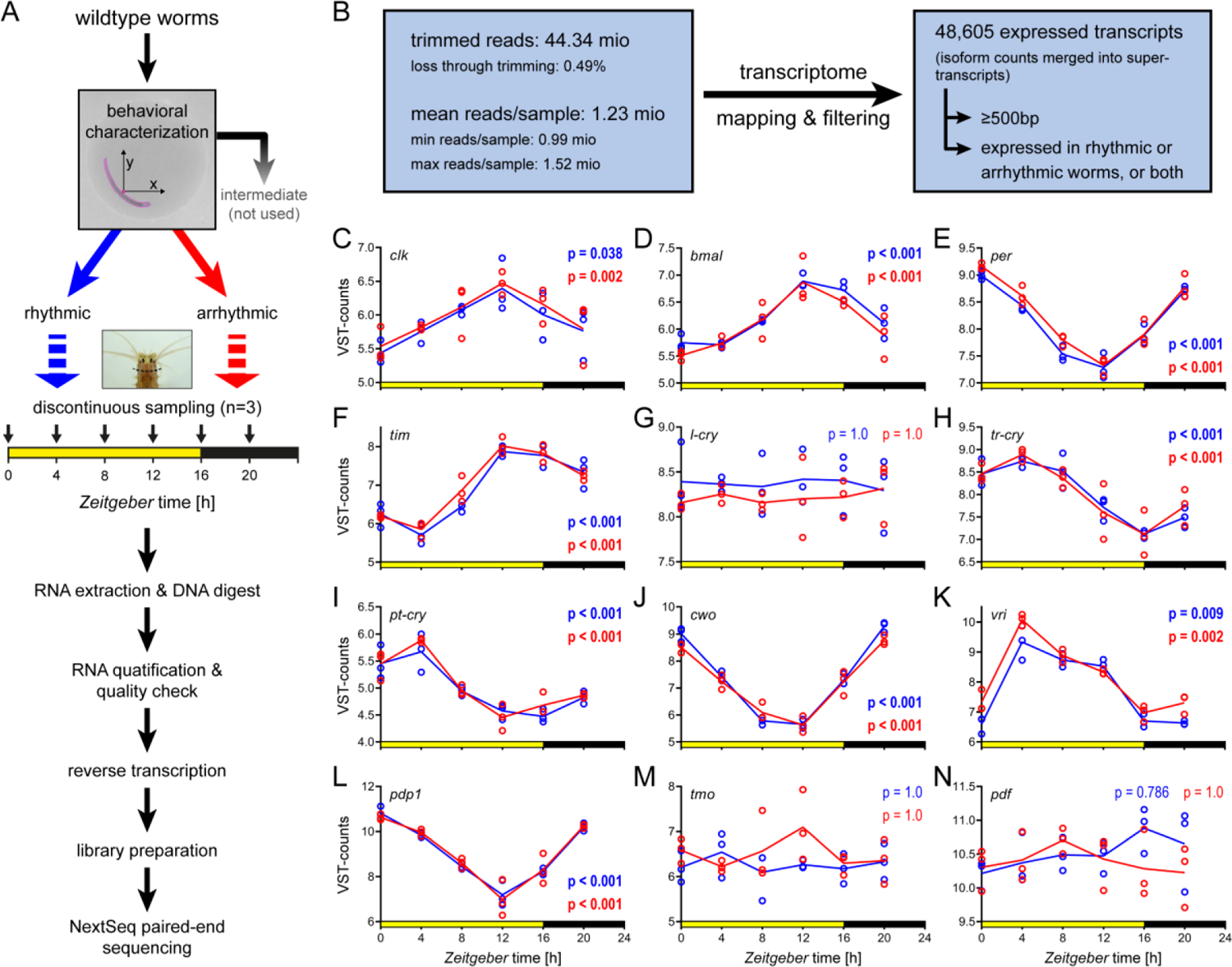
DEseq analyses reveal identical core circadian clock transcript cycling in rhythmic versus arrhythmic *Platynereis*. (A) Diel locomotor behavior of PIN strain wildtypes was characterized as rhythmic (blue) or arrhythmic (red). Worm heads were sampled discontinuously over a 16h:8h LD cycle (n=3 biological replicates (BR) per time point, 3 heads pooled/BR). Individual actograms are provided in S4 Fig. (B) Overview of the sequencing analysis pipeline. (C-N) Diel expression patterns of core circadian clock and neuropeptide transcripts. Expression is shown as VST-counts (see Methods). Bold *p*-values indicate significant 24h cycling as determined by RAIN analysis. For abbreviations of genes and detailed RAIN results see S1 Tab.

We first focused on core circadian clock transcripts (Fig 2C-N) and overall transcriptome rhythmicity. Core circadian clock transcripts of the behaviorally rhythmic worms also served as benchmark to assess if the sampling had affected transcriptome rhythmicity. Core circadian clock gene patterns (Fig 2C-L) closely mirrored previous reports [28,35,39], emphasizing reproducibility.

To further look for possible differences in transcriptomic cycling between rhythmic and arrhythmic worms, we identified transcripts with 24h cyclic expression via RAIN [40]. We detected a total of 1451 transcripts (2.99% of all expressed transcripts) with 24h cycling in rhythmic worms (n=603), arrhythmic worms (n=554), or both phenotypes (n=294) (Fig 3A and S1 Tab). Thus, the number of transcripts with significantly diel cycling was comparable between rhythmic and arrhythmic worms (Fig 3A). This suggests that behavioral arrhythmicity does not imply a generally lower rhythmicity on the level of gene expression.

**Fig 3:**
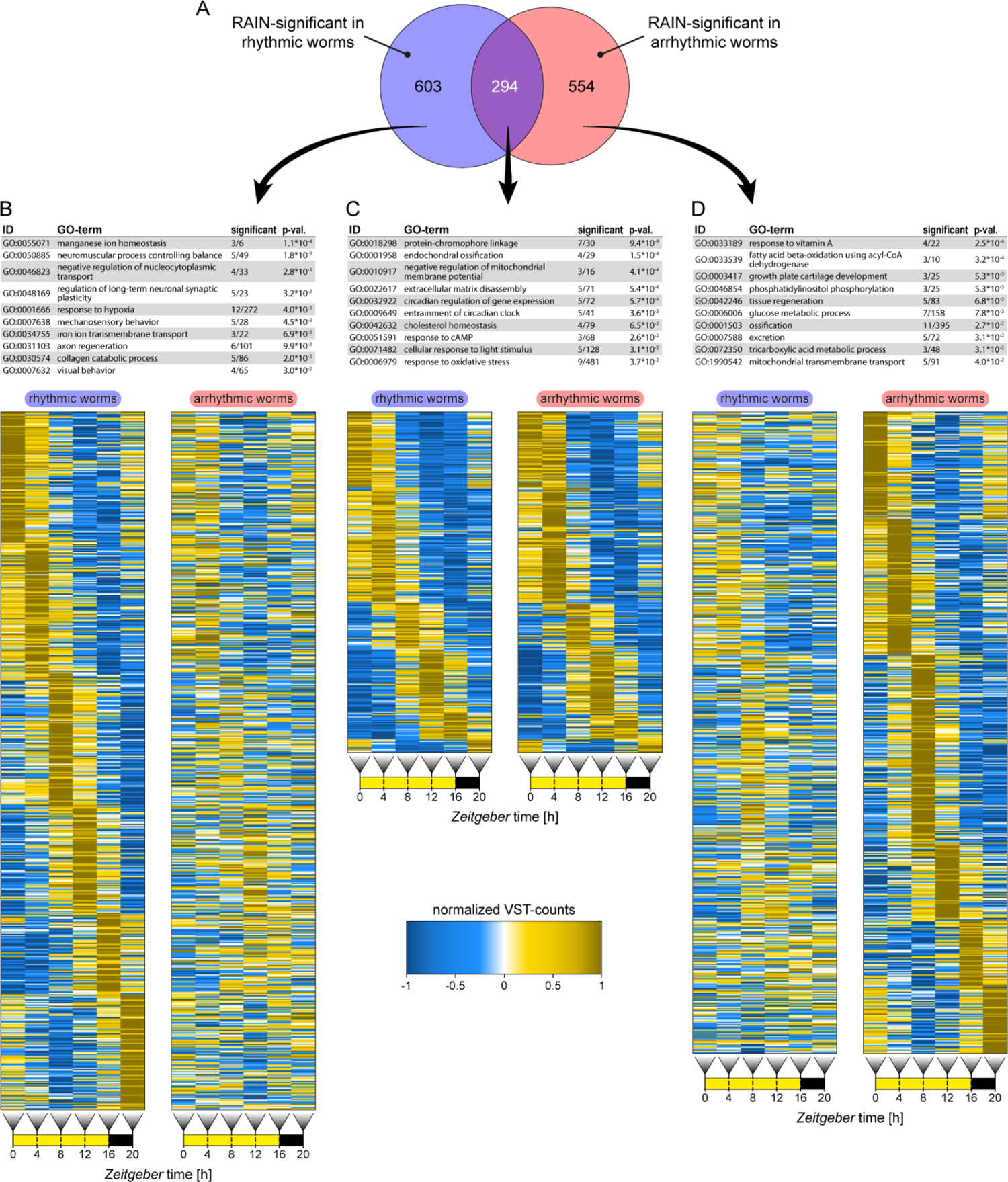
Representation of cyclic transcripts and GO-term annotations. (A) Transcript identified by RAIN analysis with significant 24 hour cycling in behaviorally rhythmic worms (blue), arrhythmic worms (red), or both (S1 Tab). (B-D) The top 10 most representative GO-terms for each group are shown, for full lists of significant terms see S2 Tab. Diel expression of transcripts significant by RAIN analysis in either group are plotted as comparative heatmaps for both groups. Heatmaps are based on mean VST-counts (see Methods) and are sorted by peak expression times. Expression is normalized for both phenotypes together, meaning amplitudes are comparable between rhythmic and arrhythmic worms, but not between different transcripts.

For the core circadian clock genes, RAIN found significant cycling in all of them for both phenotypes, except for *l-cry*, *tmo* and *pdf* (Fig 2C-N), consistent with previous results [35,39,41]. Of note, core circadian clock transcripts kinetics were basically identical between rhythmic and arrhythmic worms (Fig 2C-L). Aside from the core circadian clock transcripts, both behavioral groups showed significant common cycling of transcripts for *6-4 photolyase*, *r-opsin3*, *G_o_-opsin1* and *myocyte enhancer factor 2* (*mef2*) (S1 Tab).

### Functional categorization by Gene Ontology (GO-) term analysis suggest strong metabolic rhythmicity in behaviorally arrhythmic worms

We next categorized the cyclic transcripts for their biological functions by GO-term analysis and subsequently investigated if any terms occurred significantly more frequently in either behaviorally rhythmic or arrhythmic worms. The annotations of RAIN-significant cycling transcripts present in rhythmic, arrhythmic or both behavioral phenotypes were compared against all GO-terms of the expressed 48,605 unique transcript IDs (S2 Tab). This identified 77 GO-terms as enriched in the transcripts cycling in rhythmic worms, 29 in the transcripts cycling in arrhythmic worms, and 25 terms in the transcripts cycling in both phenotypes (Fig 3, S2 Tab).

Cycling transcripts shared by worms of both phenotypes showed enrichment in GO-terms associated in rhythmic/circadian processes such as „circadian regulation of gene expression‟ and “entrainment of circadian clock” (Fig 3C), consistent with the described indistinguishable transcript kinetics (Fig 2C-L). There were further a number of GO-terms associated with light perception like ‟protein-chromophore linkage‟ and „cellular response to light stimulus‟. Transcripts significantly cycling only in rhythmic worms showed enrichment for a variety of GO-terms with a notable accumulation of terms associated with nervous system function and behavior. Five of the top 10 most representative GO-terms were: „neuromuscular process controlling balance‟, „regulation of long-term neuronal synaptic plasticity‟, „mechanosensory behavior‟, „axon regeneration‟, visual behavior‟ (Fig 3A,B). Similar terms were not enriched in transcripts cycling only in arrhythmic worms (Fig 3D). In behaviorally arrhythmic worms, the types of significantly cycling transcripts markedly differed from those of the rhythmic worms. They showed an enrichment for GO-terms associated with several metabolic and physiological pathways (Fig 3D). This highlights that behavioral rhythmicity and metabolic/physiological rhythmicity do not necessarily correlate and can in fact be opposite. For details on the oscillation amplitudes of transcripts from the different categories please consult the supplementary material (S1 Text, S5 Fig).

In addition to these “neuronal/behavioral” versus “metabolic/physiological” terms, the GO-term „extracellular matrix disassembly‟ and other related to the extracellular space appeared prominently for both behavioral groups (Fig 3B-D). An investigation of the associated transcripts showed that many encode for extracellular peptidases of the matrix metalloproteinase (MMP) family. We next scanned all expressed transcripts for potential MMPs, followed by validation via BLASTx against the NCBI nr-database focusing on the characteristic ZnMc-peptidase and hemopexin domains [42–44]. We were thereby able to identify at least 20 putative *Platynereis* MMPs (S3 Tab). Sequence matches for these MMPs in the *Platynereis* genome (Simakov O and Arndt D, pers. comm.) further confirmed that the respective transcripts derived from individual genes. Eight of the twenty identified MMP transcripts showed RAIN-significant cycling in rhythmic worms (n=3), arrhythmic worms (n=2) or both phenotypes (n=3) (Fig 3, S6 Fig, S3 Tab).

### *Platynereis* possesses pigment-dispersing factor (PDF) and a functional PDF-receptor ortholog

In *Drosophila*, MMP1 mediates circadian axonal remodeling of clock neurons and their interactions in a system that ensures proper behavioral rhythmicity and time-of-day dependent responses of clock and behavior to external cues [33,45,46]. In flies the remodeling by MMP1 depends on the neuropeptide PDF [33,34,46]. PDF is a conserved neuropeptide that is central to the communication and output of circadian clock neurons, thereby shaping diel behavioral rhythmicity in drosophilid flies and other insects [13, 47]. Given this connection between MMP1, PDF and circadian behavioral control in *Drosophila*, and the differential cycling of several MMP transcripts occurring in our DEseq analyses, we wondered about a possible role of PDF in *Platynereis* behavioral rhythmicity. We identified *Platynereis* sequences of the PDF precursor protein (prepro-PDF) and PDF-receptor (PDFR) and performed phylogenetic analyses (Fig 4A,B, S4 Tab). This confirmed the identity of our PDF and PDFR sequences [48].

**Fig 4:**
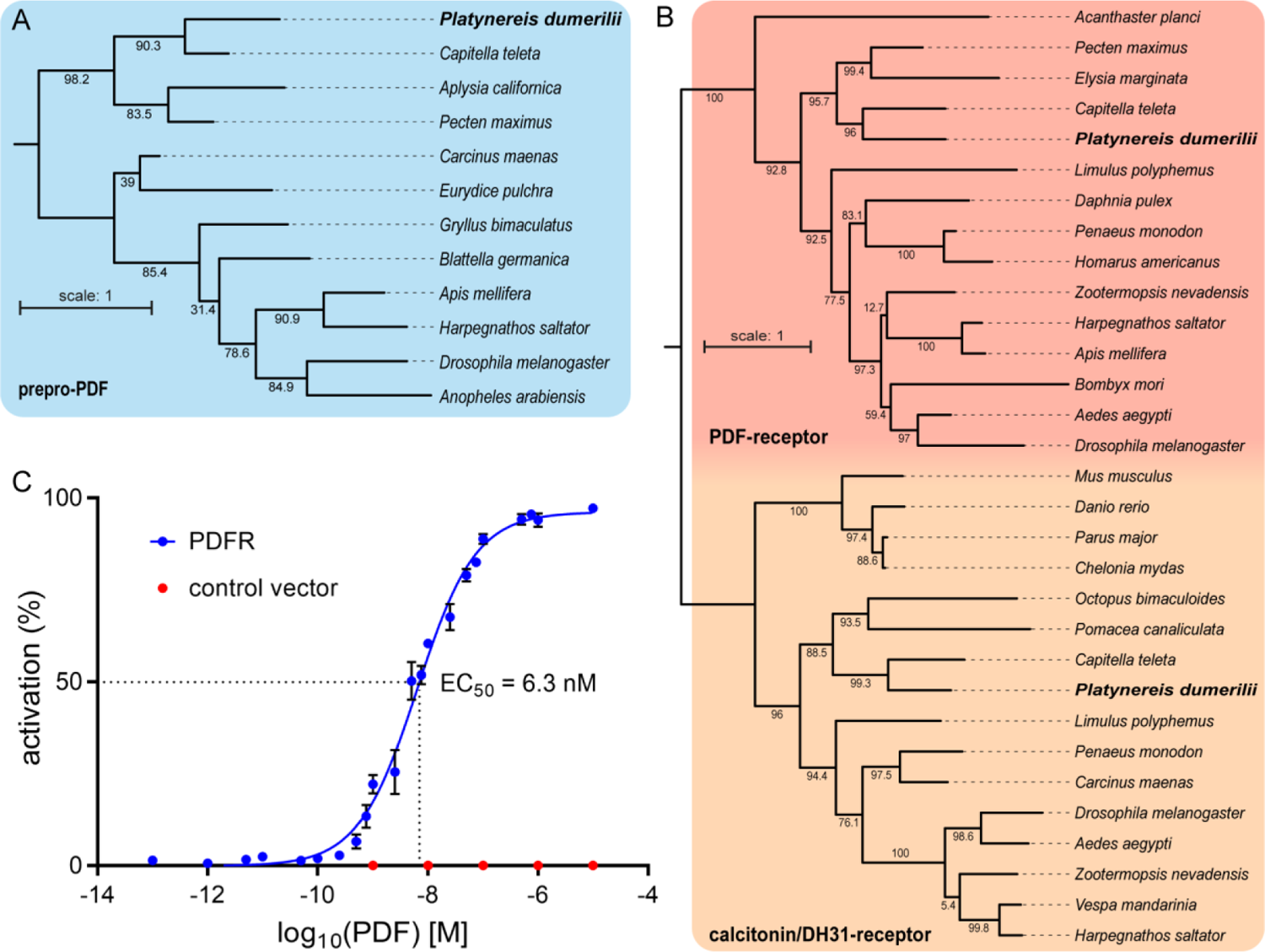
PDF and PDF-receptor phylogeny and deorphanization. Maximum-likelihood phylogenetic trees of (A) the pigment-dispersing factor precursor protein (prepro-PDF) and (B) the pigment-dispersing factor receptor (PDF-receptor), using the calcitonin-receptor and diuretic hormone 31 (DH31) receptor as outgroup. See supplement for peptide sequences used for tree construction (S4 Tab). (C) Activation curve of the *P.dumerilii* PDF-receptor by PDF (blue) with EC_50_ indicating the concentration needed for 50% receptor activation (n=10-12 per concentration). Red dots indicate activation after transfection with empty control vector (n=6 per concentration). Mean values ± SEM are shown (S5 Tab).

To validate the functionality of the PDF/PDFR-system in *Platynereis, pdfr* was expressed in CHO cells employing an *apoaequorin*/*human Gα16 subunit* reporter system [49]. The assay confirmed that in *Platynereis* PDFR can be activated by PDF at physiological concentrations (EC_50_=6.3 nM, Fig 4C).

Next, we induced mutations in the *pdf* gene of VIO wildtype worms using TALENs [50], which resulted in insertion/deletions in the 3^rd^ exon preceding the start of the mature peptide sequence (Fig 5A,B). We tested F_1_-outcrossed worms for changes in the *pdf* sequence. Worms that showed frame-shift mutations and early stop codons were again crossed against VIO wildtypes (at least 3 times). The resulting heterozygous offspring were in part incrossed to start the analyses of homozygous mutant worms, as well as further outcrossed. Thereby we established two mutant allele lines with a −14bp deletion and a +4bp insertion, respectively (Fig 5A,B). Both alleles are predicted to result in a complete failure of mature peptide generation (Fig 5B).

**Fig 5:**
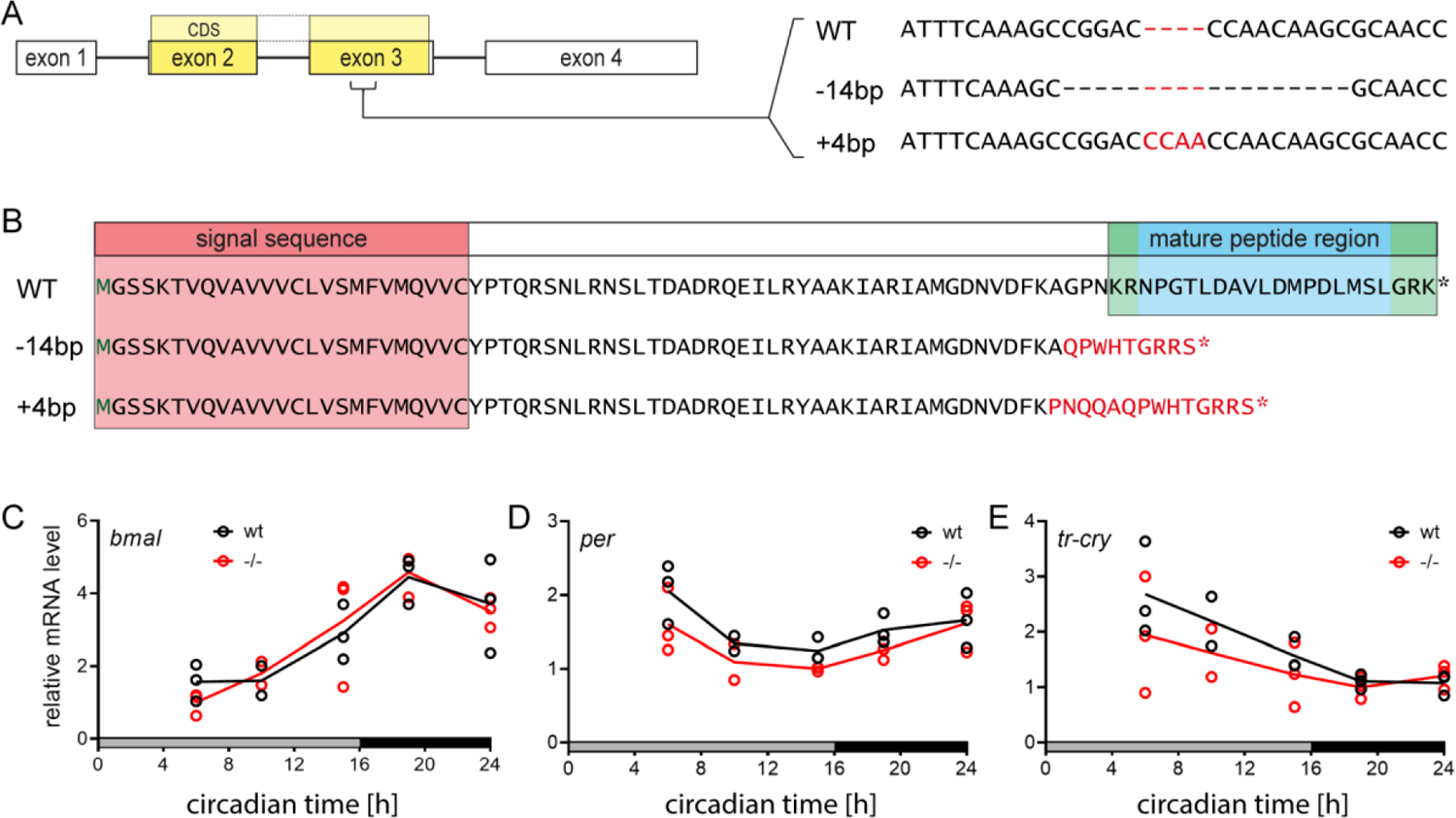
TALENs-induced *pdf* gene knockout and effects on clock genes. (A) TALENs targeting exon 3 of the *pdf* gene were used to induce frame shift mutations in the coding sequence (CDS). Two mutant alleles, a −14bp deletion and a +4bp insertion, were established. (B) Translated peptide sequence from the *pdf* wt and mutant loci. Start codon methionine (green M), signal sequence for cellular export (red background), cleavage regions (green background) and mature peptide (blue background) are indicated. Both frame shift mutations led to a premature stop codon (*) and a complete loss of the mature PDF peptide. (C) Circadian expression of the clock genes *bmal*, *per*, and *tr-cry* in *pdf* wildtypes and mutants. Samples were collected on the 2^nd^ day of DD after being cultured under LD (16h:8h). Per time point, n=3 replicates were measured (n=2 for circadian time 10). Unpaired 2-sided t-tests for each time points found no significant differences between wildtypes and mutants. Raw C_t_-values are provided in the supplement (S6 Tab).

We determined the potential effects of *pdf* loss on the circadian clock by comparing the expression of the clock genes *bmal*, *per* and *tr-cry* in *pdf* wildtypes and −14/-14 mutants. Worms initially cultured under LD (16h:8h) were transferred to DD. Clock gene expression in worm heads was analyzed by qPCR on the 2^nd^ day of DD at 5 circadian time points. Unpaired 2-side t-tests found no significant differences between wildtypes and mutants for any gene and time point (Fig 5C-E).

### *pdf* mutants show consistent increase in locomotor activity levels

To investigate the role of PDF in *Platynereis* circadian behavior, the activities of *pdf* wildtypes and mutants in a mixed VIO/PIN strain background were recorded for 4 days of LD (16h:8h) followed by 8 days of DD (Fig 6A). The behavioral characterization showed no systematic differences between mutant alleles (−14/-14, +4/+4), as well as transheterozygous worms (−14/+4), and thus data were combined (S7 Fig). *pdf* mutants consistently showed significantly elevated locomotor activity over the whole 24h cycle both in LD and DD (Fig 6B). While we initially observed a small decrease in rhythmicity under LD and DD (S8 Fig), this decrease disappeared upon further outcrossing (Fig 6, S9 Fig). In fact, the rhythmicity trend was reversed in lines that had been further outcrossed, showing an increased rhythmicity under LD, but not DD (Fig 6C,D, S9C,D Fig). As this effect was consistently maintained in more outcrossed worms, independently if backcrossed to VIO or PIN strains (Fig 6 vs. S9 Fig), we interpret the initially observed decrease in rhythmicity (S8 Fig) rather as an unspecific background effect. We can however not fully exclude that strain background might play an additional role. We believe that trusting the results from more outcrossed animals is a justified approach, and hence refer to them further below.

**Fig 6:**
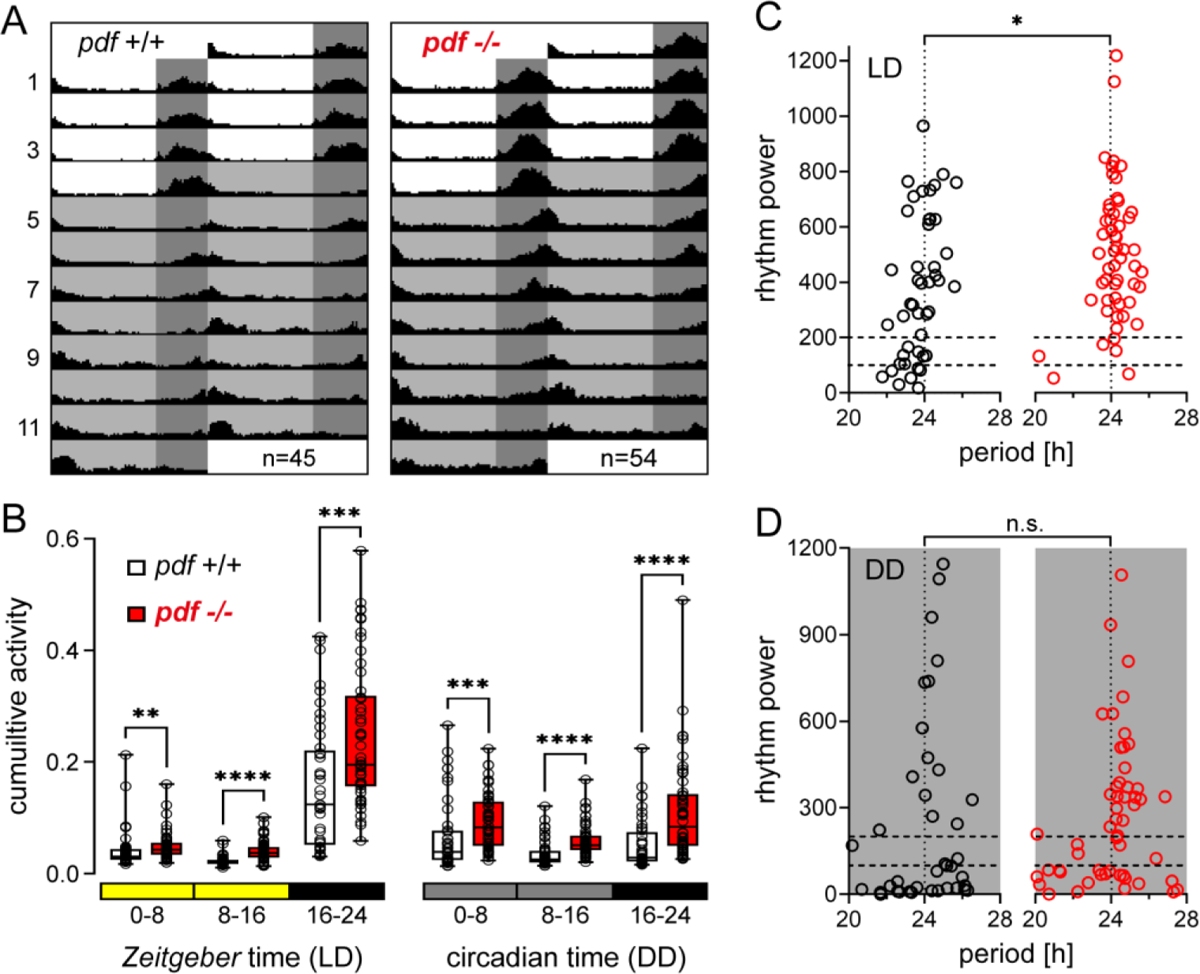
*pdf* modulates overall locomotor activity, but not rhythmicity. (A) Circadian locomotor activity of *pdf* wildtypes (black) and mutants (red) under 4 days of LD and 8 days of DD. Individual worm actograms are provided in S7A,B Fig). (B) Cumulative activity over the early day (0-8), late day (8-16) and night (16-24) in LD and DD. (C,D) Period/power of wildtype and mutant locomotor rhythms in the circadian range (20h-28h) in LD and DD determined by Lomb-Scargle periodogram. Statistical analyses (panel B-D) were performed via Mann-Whitney U-test.

The comparison of locomotor activity between LD and DD further showed that the *pdf* wildtype tends to spread out its activity more across the 24hrs under DD (S7 Tab), consistent with previous observations [35]. Under LD, part of the nocturnal worm‟s activity is thus likely suppressed by the acute light (masking). In the mutants, this might compress part of the overall increased activity into the available dark portion (S7 Tab), providing a possible explanation for the consistently observed significantly higher rhythmicity of the mutants under LD, but not DD (Fig 6C,D, S9C,D Fig).

Together, the results suggest that while PDF controls behavior in *Platynereis*, it is not a major determinant of circadian behavioral rhythmicity. This further indicates that impacts of MMPs on individual circadian rhythmicity differ from the PDF-mediated mechanism described in *Drosophila*.

## DISCUSSION

Diversity of rhythmic phenotypes, even among closely related individuals, is ubiquitous in animals. However, especially for “non-conventional” but evolutionary and ecological informative model systems investigations are still largely limited to the behavioral level [51–53]. However, it is largely unclear, how much rhythmic behavior can serve as an indicator of the overall rhythmicity of an individual, and in extent, the population. Here we uncovered circadian behavioral patterns of the marine annelid worm *Platynereis dumerilii*, which are individual-specific and persist over time. Differences in the strength of behavioral circadian rhythms were not mirrored by the transcript oscillations of all core circadian clock genes, nor in the number of overall cycling transcripts. Circadian behavioral rhythm differences were, however, mirrored by the types of transcripts showing cyclic expression: neuronal/behavioral versus metabolic/physiological transcripts correlating with behaviorally rhythmic versus arrhythmic worms, respectively.

It is noteworthy that within a (natural) population behavioral and physiological phenotypes exist as a continuum from highly rhythmic to arrhythmic, with individual-specific pattern likely emerging as the cumulative sum of various minor gene effects. The individual diversity in worm diel phenotypes is remarkable, especially given that many of the investigated worms were siblings and were raised together in the same culture boxes. The high consistency of behavioral patterns over time illustrates that they are not just behavioral snapshots of phenotypic plasticity, but individual-specific characteristics. Similarly consistent individual behavior patterns have been described in several other species [25, 54]. In a striking example, Bierbach et al. raised genetically identical freshwater fish under near-identical conditions, but nevertheless observed individual-specific behavior patterns that persisted over several rounds of investigations, suggesting environmental influences possibly via epigenetic modifications [55]. Such modifications might also explain the similar diversity of individual rhythmicity in *Platynereis* highly inbred versus non-inbred strains and as such provides an interesting level of plasticity across a population.

### Ecological benefits of rhythmic diversity – beyond the individual

Control of diel rhythmicity by circadian clocks is a major factor shaping a species‟ adaptations to its cyclic environment and can thus strongly affect fitness [10,11,56,57]. According to the classical view, endogenous clocks help to anticipate cycles and optimize timing to maximize resource usage and minimize predation risk. In this context, the diversity of rhythmic phenotypes observed in *P. dumerilii* independently of core circadian clock transcript-cycling, might appear initially contradictory. It will however allow a species to inhabit a broader range of microhabitats and survive under relatively sudden ecological changes.

As we found no clear connection between *Platynereis* rhythmic diversity and a specific genetic background (strain), we reason that early environmental factors will likely play a role. First, rhythmic diversity could be a form of bet-hedging, likely via epigenetic variations that are induced either maternally or by stochastic environmental conditions experienced during a very early life phase [55,58,59]. Bet-hedging has been stated as a potential adaptation of seasonal life cycles to interannual variability [60], but has so far not been discussed in the context of diel rhythmicity. Second, rhythmic diversity in *Platynereis* could be established upon settlement of the planktonic larvae on the sea floor [27]. The conditions encountered at this time may lead to an „imprinting‟ of a phenotype that seems most suitable for the respective microhabitat.

Diversity of diel rhythms and other phenotypic traits may be especially important in marine habitats. Various marine species including *Platynereis* practice broadcast spawning where gametes are dispersed by currents as part of the plankton, before the larvae settle on the seafloor with limited influence on their final settlement place. By creating offspring with a variety of phenotypes, bet-hedging increases the chances of at least some individuals being well-adapted, especially when environmental conditions are variable in time and/or space [61, 62] as it is the case in the coastal waters inhabited by *Platynereis* [28]. Phenotypic diversity can further reduce intraspecific competition due to a diversification of resource usage and spatial/temporal niches [63, 64].

Individual diversity of rhythmicity (and in general) is an important factor to consider when investigating tolerance to environmental conditions. Tolerance ranges are commonly reported as averages or 50% lethality limits for entire populations or species. Our transcriptome analysis shows that rhythmic/arrhythmic individual phenotypes are associated with different physiological adjustments, and physiological patterns in general can be individual-specific [62, 65]. This should result in individual-specific tolerance ranges for environmental conditions. Furthermore, patterns of behavioral rhythmicity can directly determine the environmental conditions and cycles experienced by an animal [66]. Thus, considering individual diversity in experimental planning (using multiple strains) and discussing it not as „background noise‟ but as a biological feature can advance our understanding of population- and species-level adaptability to changing environmental conditions.

### Transcript cycling patterns suggest opposite rhythmicity strength in behavior versus physiology

An unanticipated finding was that wildtype worms characterized as rhythmic or arrhythmic showed identical diel expression patterns for the circadian clock genes. The few transcriptomic investigations of individual rhythmicity existing so far found correlations between diel activity pattern and different circadian clock components in forager versus nurse ants [19], as well as phenotype-specific expression of cryptochromes in zebrafish [54]. In *Drosophila*, while a transcriptome comparison of early/late emerging fly chronotypes from different strains found no difference in clock genes [67], phenotype differences in diel activity have been linked to the clock gene *pdp*1 as well as the output genes *pdf* and *timeout* [18]. Our results indicate that in *Platynereis* individual-specific rhythmic differences are not driven by circadian clock transcript cycling, nor by the action of PDF.

Instead they correlate with pathways downstream of the circadian clock, thus contrasting with reports from other species [19,54,67]. It should be noted that we used whole heads while most *Drosophila* studies looked at specific brain regions or neurons. Hence, phenotype-specific clock gene patterns on the level of individual neurons cannot be fully excluded, although such patterns have so far not been described in *Platynereis* [35].

The fact that worms that are behaviorally clearly rhythmic or arrhythmic show much more nuanced differences in gene expression cycling further emphasizes that rhythmicity monitored only via behavioral recordings or through a limited number of candidate genes (e.g. core circadian clock genes) is not representative of overall organismic rhythmicity. This is especially highlighted by the considerable number of transcripts cycling exclusively in behaviorally arrhythmic worms. As physiology can – much like behavior – be very much individual-specific [65], our results suggest that chronobiological phenotype investigations should routinely consider multiple circadian clock outputs from different levels of biological organization.

The enriched GO-terms associated with transcripts only cycling in either rhythmic or arrhythmic worms likely reflect the differences in the respective lifestyles. For behaviorally rhythmic worms, the cycling of transcripts linked to behavior as well as neuronal plasticity might indicate a more explorative lifestyle with regular foraging excursions. This could be beneficial in food-limited habitats. In contrast, the enrichment of metabolism/physiology-related GO-terms in behaviorally arrhythmic worms would be beneficial in habitats where food is readily available (e.g. the surface of a seagrass leaf). Under these conditions, no major excursions are necessary and a rhythmic optimization of physiology could improve nutrient processing, growth and overall fitness. The enhanced metabolic/physiological rhythmicity in behaviorally arrhythmic worms again illustrates that behavioral outputs alone may not be representative and that integrative investigations of organismal rhythmicity can be highly beneficial in revealing unexpected rhythmic complexity in any species.

## Conclusions

We identify consistent circadian rhythmic diversity among *Platynereis dumerilii* strains and individuals, correlated with previously unknown molecular features. Individual *Platynereis* worms can be discriminated based on their reproducible circadian locomotor patterns. The behavioral rhythmic phenotypes are independent of circadian clock transcript cycling and, in contrast to *Drosophila* [68], genetic ablation of the neuropeptide PDF has no effect on *Platynereis* circadian rhythmicity. Our results argue that circadian rhythmicity cannot be defined through stereotypic behavioral phenotypes or indicator genes alone, but that it is the sum of various processes acting on different levels of organization. Most notably, our observation of enhanced diel cycling of metabolic/physiological transcripts in behaviorally arrhythmic worms highlights the complex relationship between the circadian clock and its behavioral and physiological outputs.

Investigating the factors that shape phenotypic and rhythmic diversity in an integrative way will help to understand how animals adapt to specific habitats and how these factors can contribute to overall population fitness. It highlights the importance of treating biological variation not as „background noise‟, but as an additional aspect of biodiversity that needs to be considered in both the mechanistic and the ecological context [24, 69].

## MATERIALS & METHODS

### Experimental model & subject details

We performed all investigations on the marine annelid *Platynereis dumerilii* (Audouin & Milne Edwards, 1833). The worms used belonged to 3 wildtype strains, which all originated from worms collected in the Gulf of Naples, Italy (40.7°N, 14.2E). The PIN and VIO strains have been in culture for ∼70 years and have been incrossed at least 10 times [30, 37]. The NAP strain was established in 2009 and has been kept as non-incrossed closed culture. For all experiments we used only immature/premature worms (age: 3-7 months, min. length: ∼2.5 cm) that did not show any signs of metamorphic maturation to the adult spawning form (epitoke).

### Worm culture conditions

All worms used in the described experiments were cultured at the Max Perutz Labs (Vienna, Austria) in a 1:1 mix of natural seawater collected in the German bight by the Alfred-Wegener-Institute (Bremerhaven, Germany) and artificial seawater (Meersalz CLASSIC, Tropic Marine AG, Switzerland). The water was pumped through 1 µm filters before use and was exchanged every 1-2 weeks. Worms were kept in transparent plastic boxes (30×20 cm, 100-150 indv./box) in temperature controlled rooms at 19.5±1.0°C under a rectangular 16h:8h light/dark-cycle (LD) with fluorescent tubes (4.78*10^14^ photons*cm^-2^*s^-1^, Master TL5 HO 54W/865, Philips, Netherlands, S1A,C Fig). To account for *Platynereis*‟ lunar reproductive cycle [35,70,71], we simulated a moonlight cycle by an LED lamp (Warm White 2700K A60 9W, Philips, Netherlands) that was switched on for 8 nights centered on the outdoor full moon times (see [36] for details). Larvae worms were fed a mix of living *Tetraselmis* algae and *Spirulina* (Organic Spirulina Powder, Micro Ingredients, USA) 2-3 times a week until they started forming their living tubes. Thereafter, worms were fed shredded spinach and powdered fish flakes (TetraMin, Tetra GmbH, Germany) (once a week, each). Under these conditions, *Platynereis* has a generation time of min. 3 months.

### Recording and analysis of worm locomotor behavior

We investigated worm locomotor activity in light-tight chambers developed in cooperation with loopbio GmbH, Austria [28, 72]. The worms were last fed (spinach) 4 days before the start of the investigation. Worms were always taken from culture boxes of similar animal densities and age to avoid housing effect on behavior. To prevent potential behavioral changes associated with the simulated lunar cycle (see above), we conducted all recordings in the 2 weeks around new moon. For recording, the worms were transferred to 5×5 well plates and were filmed at 15 fps using infrared-illumination and a camera equipped with a long-pass IR-filter (acA2040-90um, Basler AG, Germany). A custom-build LED-array was used to create a 16h:8h LD-cycle with natural spectral composition (1.40*10^15^ photons*cm^-^²*s^-1^, Marine Breeding Systems GmbH, Switzerland, S1B,C Fig). All recordings were done at culture temperature and diel oscillations were ∼1°C in LD and <0.5°C in constant darkness (DD). For data analysis, we excluded worms that matured during or within one week after the recording, as worm behavior is strongly altered during the maturation phase. Data of worms that crawled out of the well and could no longer be recorded were omitted, if less than 3 days of data were available for the respective recoding phase (LD or DD). Sex determination in a limited number of wildtype worms showed no sex-specific differences in locomotor behavior (S2 Fig) and hence data of males and females were pooled.

We quantified locomotor activity by determining x/y-coordinates of the center of the detected worm shape (Fig 2A) and measuring the distance moved between frames with the Motif automated tracking software (loopbio GmbH, Austria, see [28] for details). We used a 1-min moving average to reduce noise. Period and power of diel rhythmicity was determined via Lomb-Scargle periodograms with the ActogramJ plug-in in ImageJ/Fiji [73, 74]. Analysis focused on the 20h-28h range of circadian rhythmicity. Overall activity was binned into morning (ZT0-8), afternoon (ZT8-16) and night (ZT16-24) for the LD phase, and into subjective morning (circadian time, CT0-8), subjective afternoon (CT8-16) and subjective night (CT16-24) for the DD phase with all days of the respective phases being pooled.

We compared diel rhythmicity power between groups, as well as overall activity between time bins (morning, afternoon, night) and between groups via unpaired 2-side t-test or via One-way ANOVA with Holm-Sidak *post-hoc* test. If data were not normally distributed and/or homoscedasticity of variances was not given, we replaced the t-test with a Mann-Whitney U-test and the One-way ANOVA with a Welch‟s ANOVA with Dunnett-T3 *post-hoc* test or a Kruskal-Wallis ANOVA on Ranks with Dunn‟s *post-hoc* test. Statistical analyses were carried out in GraphPad Prism (GraphPad Software Inc., USA).

The number of recording days in LD or DD depended on the specific investigation. For the comparison of the PIN, NAP and VIO wildtype strains (Fig 1), we exposed the worms to 4 days of LD and DD, respectively. We recorded n=25 PIN worms, n=21 NAP worms and n=38 VIO worms (Fig 1, S2 Fig). PIN, NAP and VIO worms for the comparison originated from 2 mating batches, each. Repetition runs were done with 12 PIN worms, 5 NAP worms and 13 VIO worms (Fig 1E-G and S3 Fig) and the repeated worms were kept separately in smaller boxes after the initial recording to avoid the confusion of individuals.

To behaviorally characterized worms for RNASeq analysis (Fig 2, described below) we exposed them to 3 days of LD and DD, respectively. Over the course of 6 months, we performed 13 behavioral recordings using a total of 325 PIN wildtype worms from 5 different mating batches.

Worms in the comparison of *pdf* mutants and wildtypes (Fig 6) had a mixed VIO/PIN background and experienced 4 LD days followed by 4, 5 or 8 DD days. The fractions of worms experiencing a given number of DD days did not differ between wildtypes and mutants, meaning that datasets are directly comparable. We performed 5 recordings using n=45 *pdf* wildtypes from 5 mating batches and n=54 *pdf* mutants from 6 mating batches (−14/-14 n=16, +4/+4 n=31, −14/+4 n=7). As the mutants showed no within-strain differences among genotypes (S4 Fig), all mutant genotypes were pooled. Details on the recordings of *pdf* worms in the initial VIO strain background (S8 Fig) and after later being backcrossed into the VIO strain (S9 Fig) are provided in the respective supplemental material. The *pdf* genotypes of all mutants and the associated wildtype worms were verified at least 1 month before the behavior recording or directly afterwards (described below).

Actograms of individual worms used in strain comparison, characterization for RNASeq, and *pdf* wildtype/mutant comparison are provided in the supplement (S2,4,7 Fig).

### Rhythmic characterization of wildtype worms and sampling for RNASeq analysis

PIN wildtype worms whose locomotor activity had previously been recorded (3 days LD & DD, see above) were first grouped based on their behavioral rhythmicity. We characterized worms as rhythmic (n=102) or arrhythmic (n=75) based on Lomb-Scargle periodogram analysis and visual inspection of actograms. For the characterization, rhythmicity in DD was considered more important than in LD. Worm that were neither clearly rhythmic nor arrhythmic were considered „intermediate‟ and were not used further (n=148 incl. maturing worms). While the relative abundance of phenotypes did partially differ between recordings, all included rhythmic and arrhythmic worms. Actograms of all individual worms including their rhythmic characterizations are provided in S4 Fig.

We sorted characterized worms to culture boxes based on rhythmicity phenotype and kept them under standard conditions (LD 16h:8h + lunar cycle) until sampling at the next new moon (i.e. ∼1 month after the behavioral characterization). Samples for RNASeq analysis were collected at ZT0 (lights on), ZT4, ZT8, ZT16 (lights off) and ZT20 (Fig 2A). Due to the work-intensive characterization procedure and the loss of characterized worms due to maturation, we could not collect all samples for the diel time series „in one go‟, but sampled discontinuously over several months while making sure that replicates of the same rhythmic phenotype/time point were not collected simultaneously. Worm age at the time of sampling was 4-7 months. We last fed the worms 5 days before sampling (spinach) and 2 days before sampling we transferred them to sterile filtered seawater containing 0.063 mg/mL streptomycin-sulfate and 0.250 mg/mL ampicillin to minimize bacterial contaminations. Worm heads were sampled by anesthetizing animals in 7.5% (w/v) MgCl_2_ mixed 1:1 with sterile filtered seawater for 5 minutes. Heads were cut before the first pair of parapodia under a dissection microscope before being fixed in liquid nitrogen with 3 heads pooled per replicate and samplings at ZT16 and TZ20 conducted under dim red light. For each time point, we collected 3 replicates of rhythmic and arrhythmic worms, respectively. During sampling, worms were also visually inspected for differences between rhythmic phenotypes (body size/shape/coloration, parasites), but none were found. RNA was extracted as described above and was stored at −80°C.

### Processing of RNASeq data

Quality control, reverse transcription and library preparation of rhythmic/arrhythmic samples were performed by the VBCF Next-Generation Sequencing facility services according to standard procedures (VBCF-NGS, Vienna Biocenter Campus, https://www.viennabiocenter.org/vbcf/next-generation-sequencing). Samples were run on a NextSeq550 System (Illumina, USA) with the High Output-kit as 75bp paired-end reads and resulted in a total of 445,553,522 raw reads (sample range: 9,973,579-15,234,688 reads).

We trimmed/filtered the raw reads from NextSeq sequencing using cutadapt (--nextseq-trim=20 -m 35 --pair-filter=any) with adapters being remove in the process [75]. This reduced read numbers to 443,393,107 reads (0.48% loss). Read quality was checked before and after trimming using FastQC [76]. Trimmed reads were mapped against a *Platynereis* reference transcriptome generated from regenerated tail pieces (Fig 2B). Trimmed-read fastq-files and FastQC-reports as well as transcript sequences of the reference transcriptome were submitted to the Dryad online repository. We chose this tail transcriptome, because a BUSCO analysis [77] showed that it contained a higher percentage of near-universal single-copy orthologs (98.9% complete) compared to a previously published transcriptome generated from worm heads [38]. To ensure that no important transcripts were lost due to tissue-dependent expression, we manually verified the inclusion of known clock genes, opsins, neuropeptides and other potential genes of interest. The respective transcripts were identified using internal sequence resources and NCBI, and were verified via BLASTx against the nr-database. If sequences from existing *Platynereis* resources covered larger gene stretches than the respective transcript, we replaced the sequences in the transcriptome or merged them. Trimmed reads were mapped to the reference transcriptome using salmon tool [78] (standard parameters incl. multi-mapping, library type: ISR). To exclude transcripts that did not receive any reads during mapping, we removed those transcripts from the expression matrix that had no observed expression in at least 2 time points in the rhythmic time as well as the arrhythmic samples. We further excluded all transcript <500bp to avoid fragments that could not be reliably annotated, thereby obtaining a total of 48,605 expressed transcripts for differential expression by sequencing (DEseq) analyses (Fig 2B and S1 Tab).

### Identification of cycling transcripts with RAIN

To identify cyclic changes in transcript expression patterns, we used the R-package „RAIN‟, which employs a non-parametric approach to identify cycling independent of waveform [40]. As RAIN assumes homoscedasticity, raw count data were transformed via the *VarianceStabilizingTransformation* function of the DESeq2 package and resulting VST-counts were analyzed [79]. Cycling was investigated separately for rhythmic and arrhythmic samples for a 24h period with RAIN standard settings (peak.border=c(0.3,0.7)). We could not test for other periods due to the discontinuous sample collection, which would have disrupted non-24h cycles (see above). Resulting *p*-values were false-discovery-rate corrected via the *p.adjust()* function in R according to the Benjamini-Hochberg procedure [80] with *p*-values <0.05 considered significant (Fig 3A and S1 Tab).

For visualization, we normalized VST-counts of RAIN-significant transcripts to a scale of −1 to 1 and plotted them in heatmaps using the R-packages *gplots* [81] and *RColorBrewer* [82]. Transcripts with cycling in rhythmic worms, arrhythmic worms, or both were plotted separately, but always for both phenotypes (Fig 3B-D). The normalization means that expression amplitudes of a given transcript are comparable between rhythmic and arrhythmic worms, but not between different transcripts or groups (rhythmic, arrhythmic, both).

### GO-term annotation and enrichment analysis

We next performed Gene Ontology (GO) analysis to determine the enrichment of biological processes among transcripts with RAIN-significant cycling. For this, transcripts were assigned GO-terms using the Trinotate protocol [83]. Briefly, we used TransDecoder (https://github.com/TransDecoder/TransDecoder) to identify the most likely longest open reading frame (ORF) peptide candidates and performed a BLASTx or BLASTp search of the transcripts or peptide candidates against the SwissProt database (UniRef90) [84, 85]. Additionally, we used the *hmmscan* function from HMMER (v3.3.1, http://hmmher.org/) to get functional information from protein domains by searching the Pfam database [86]. For analysis, we pooled the GO-terms hits that had e-values <1*10^-3^ in any of the annotation approaches. GO-terms could be assigned to 19,096 out of the 48,605 expressed transcripts, and to 334, 273 and 176 of the transcripts cycling in rhythmic, arrhythmic and both phenotypes, respectively.

We determined GO-term enrichment for cycling transcripts using the R-package „topGO‟ [87], using Fisher‟s exact test and the „elim‟ algorithm [87, 88]. The elim algorithm avoids redundant scoring of more general GO-terms and thereby gives higher significance to more specific terms than the „classic‟ algorithm. GO-terms were considered significantly enriched, if they had a *p*-value ≤0.05 and were represented by at least 3 transcripts (Fig 3B-D).

### Description of *Platynereis* pigment-dispersing factor (*pdf*) and receptor (*pdfr*) sequences

An initial *Platynereis pdf* gene fragment was identified from Expressed Sequence Tags (ESTs) [89]. The resulting 567bp fragment was subsequently extended to 804bp, which included the complete coding sequence (CDS) of the prepro-peptide (accession no°: GU322426.1).

A first fragment of the *Platynereis* PDF-receptor (*pdfr*) sequence was obtained by degenerated PCR and subsequently extended by RACE-PCR [90]. A 1604bp sequence containing the full CDS was cloned into a pGEM^®^-T Easy Vector (Promega, USA) and sequenced (accession no°: OL606759). A draft genome of *P.dumerilii* (Simakov et al. *in prep.*) was searched for possible gene duplicates of *pdf* using a hidden Markov model, but no further copies were identified. For the construction of phylogenetic trees for prepro-PDF and PDF-receptor (Fig 4A,B), we collected sequences from NCBI via PSI-BLAST (3 iterations), aligned them with MUSCLE [91] and then constructed the trees using the IQ-TREE online tool with default settings [92]. Sequences used for tree construction are provided in the supplement (S4 Tab).

### Deorphanization of pigment-dispersing factor receptor

To validate that the identified *pdfr* sequence encodes a functional receptor activated by PDF, a fragment of the *pdfr* gene including the full CDS was amplified by PCR and cloned into the HindIII/XbaI sites of a pCDNA3.1(+) cloning vector. We assessed receptor activation as previously described [49]. Briefly, *pdfr-*transfected Chinese Hamster Ovary (CHO) cells expressing *apoaequorin* and the *human Gα16 subunit* were incubated with 10^-13^-10^-4^ M of synthetic amidated PDF with n=10-12 measurements per concentration. PDF was provided by the Mass Spectrometry Facility of the Institute for Molecular Pathology, Vienna; and assessed by the Chemistry Department of the University of Vienna for purity using matrix-assisted laser desorption/ionization time-of-flight mass spectrometry (MALDI-TOF-MS). We recorded calcium responses for 30 sec using a Mithras LB 940 luminometer (Berthold Technologies, Germany). After these ligand-stimulated calcium measurements, Triton X-100 (0.1%) was added to each sample well to obtain a measure of the maximum calcium response and to calculate dose responses. Percentage activation values were plotted, with cells transfected with an empty pCDNA3.1(+) vector serving as negative controls with n=6 per concentration (Fig 4C and S5 Tab). The half-maximal effective concentration (EC_50_) was calculated using a computerized nonlinear regression analysis with a sigmoidal dose–response equation in GraphPad Prism.

### Generation of *pdf*-mutant strain using TALENs

To achieve a knockout of the *Platynereis pdf* gene, we followed a reverse genetic approach using TALENs (transcription activator-like effector nucleases) as previously described [50]. Repeat-variable di-residue (RVD) sequence of TALENs were designed based on the previously identified *pdf* sequence (accession nr: GU322426.1) to target a site in the 3^rd^ exon preceding the start of the mature peptide sequence. VIO strain worm zygotes 1 hour post fertilization were digested with proteinase K (23 sec) and were then injected with 10 µL injection solution, which consistent of 200 ng/µL of each TALEN, 8 µL RNAse-free H_2_O and 2 µL TRITC-dextrane (3% in 0.2 M KCl). Before injection, the solution was filtered through 0.45 mm PVDF centrifugal filters (Ultrafree-MC-HV, Merck Millipore, USA) [50].

Once grown to maturity, we outcrossed injected worms against VIO wildtype worms, and the resulting offspring were genotyped to identify changes in the PCR-product length of the *pdf* locus using the HotSHOT-method [93] and primers covering the TALENs cleavage site. PCR products were run on agarose gels [94]. When changes in length compared to the wildtype were found, the bands were cut out and sub-cloned according to manufacturer protocols into either a pjet1.2 blunt vector (CloneJET^TM^ PCR Cloning Kit, Thermo Fisher Scientific, USA) or a pGEM^®^-T Easy Vector (pGEM^®^-T Easy Vector Systems Kit, Promega, USA) depending on the Taq-polymerase used for PCR (blunt vs. sticky ends). The vectors were used to transform competent *E. coli* cells (XL1-blue, raised by lab technician), of which individual colonies were picked. Success of the transformation was verified by PCR followed by sequencing of the insert to identify base deletions/insertions [94]. We thereby established a −14bp deletion mutant strain, a +4bp insertion mutant strain, as well as a corresponding wildtype strain (Fig 5A). The mutations resulted in early stop codons and a complete loss of the mature peptide sequence (Fig 5B). The mutants were initially crossed against VIO wildtypes at least 3 times before later also being outcrossed against PIN wildtypes to create a mixed VIO/PIN background. A rhythmic phenotype (reduced rhythmicity) observed in initial VIO strain *pdf* −14/-14 and −14/+4 mutants (S8 Fig) resembled *pdf* mutants phenotypes in *Drosophila* [13]. However, the phenotype instantly disappeared after the outcross against the PIN strain (Fig 6) and could not be recovered by backcrossing against VIO worms (S9 Fig), thus strongly indicating an off-target effect in the initial VIO mutants that was unrelated to the knockout of *pdf*.

### qPCR measurements

After entrainment to a 16h:8h LD cycle, we sampled worm heads of *pdf* wildtypes and −14/-14 mutants (VIO background) on the 2^nd^ day of constant darkness at circadian time point 6 (CT6), CT10, CT15, CT19 and CT24 as described above for RNASeq analysis. Animals were only briefly exposed to light during dissection. At each time point, 3 replicates of mutant and wildtype worms were collected with 5 heads pooled per replicate (only 2 replicates for CT10). For RNA extraction, we homogenized samples in 350 µL RNAzol® (Sigma-Aldrich, USA) with a TissueLyser (Qiagen, Netherlands) for 2 min at 30Hz and extracted RNA using the Direct-Zol RNA MiniPrep kit (Zymo Research, USA) with on-column DNA digest according to the manufacturer protocol. RNA was eluded in 25µL Nuclease-free H_2_O and was stored at −80°C. We spectrometrically determined RNA concentration and purity (Nanodrop 2000, Thermo Fisher Scientific, USA). 1 µg RNA was reverse transcribed to cDNA (LunaScript RT SuperMix, New England BioLabs, USA).

We measured the expression of the circadian clock genes *brain and muscle Arnt-like protein* (*bmal*), *period* (*per*) and *tr-cryptochrome* (*tr-cry* aka *mammalian-type cry*) via real-time qPCR (StepOnePlus^TM^, Applied Biosystems, USA) using previously established primers [28] and *cell division cycle 5* (*cdc5*) acting as a reference for normalization [38] (Fig 5C-E and S6 Tab). All samples were measured as technical duplicates.

We normalized raw C_t_-values of the clock genes *bmal*, *per* and *tr-cry* according to the 2^-ΔΔCT^ method [95] using the gene *cdcd5* as a reference (Fig 5C-E). Expression stability of *cdc5* has previously been verified [38]. Raw C_t_-values are provided in the supplement (S6 Tab). Differences between worm groups were analyzed via unpaired 2-sided t-tests for each time point and with correction for the testing of multiple time points. All statistical tests were performed using GraphPad Prism.

### Resources (consumables, animals, cell lines, sequences, software) & deposited data

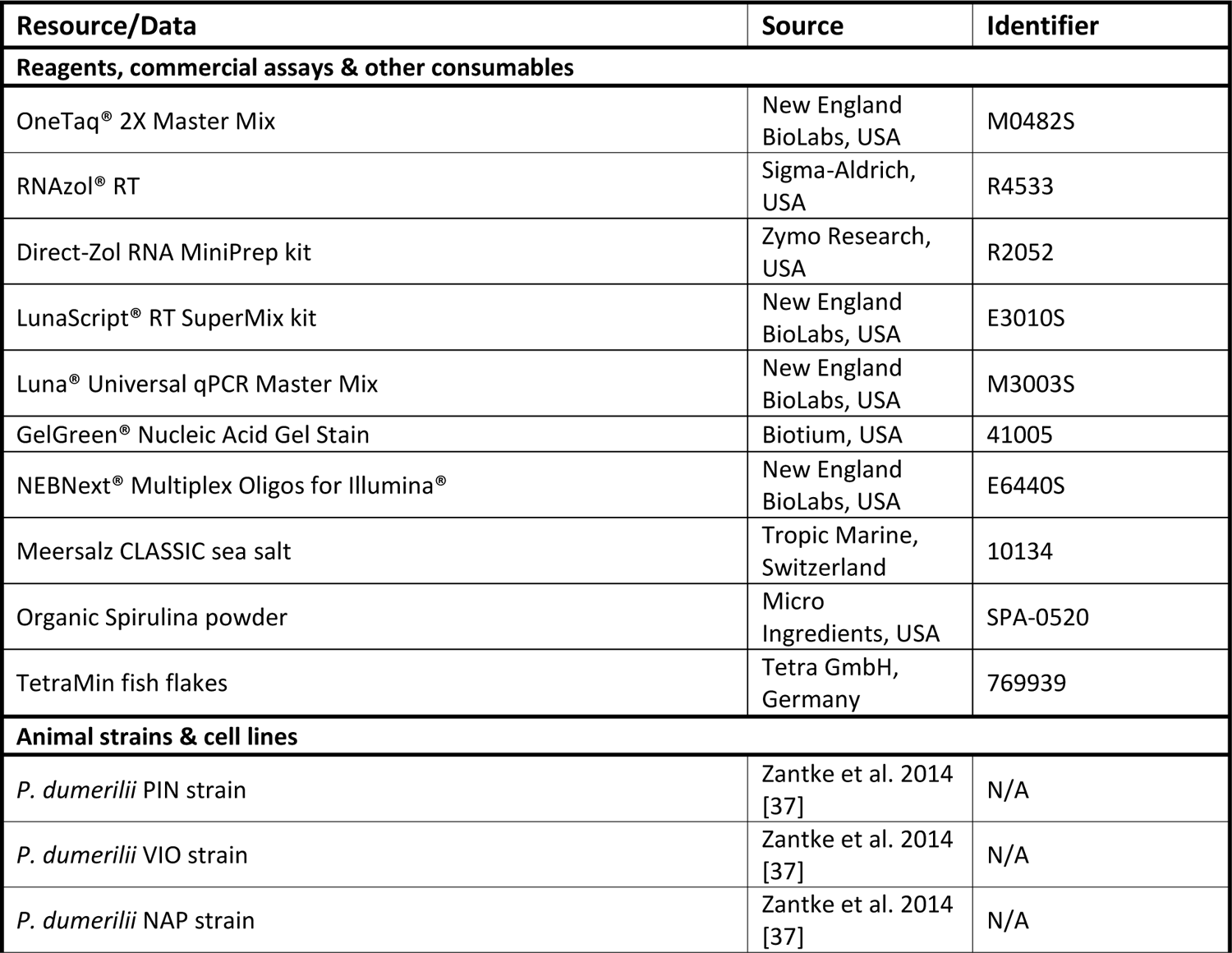
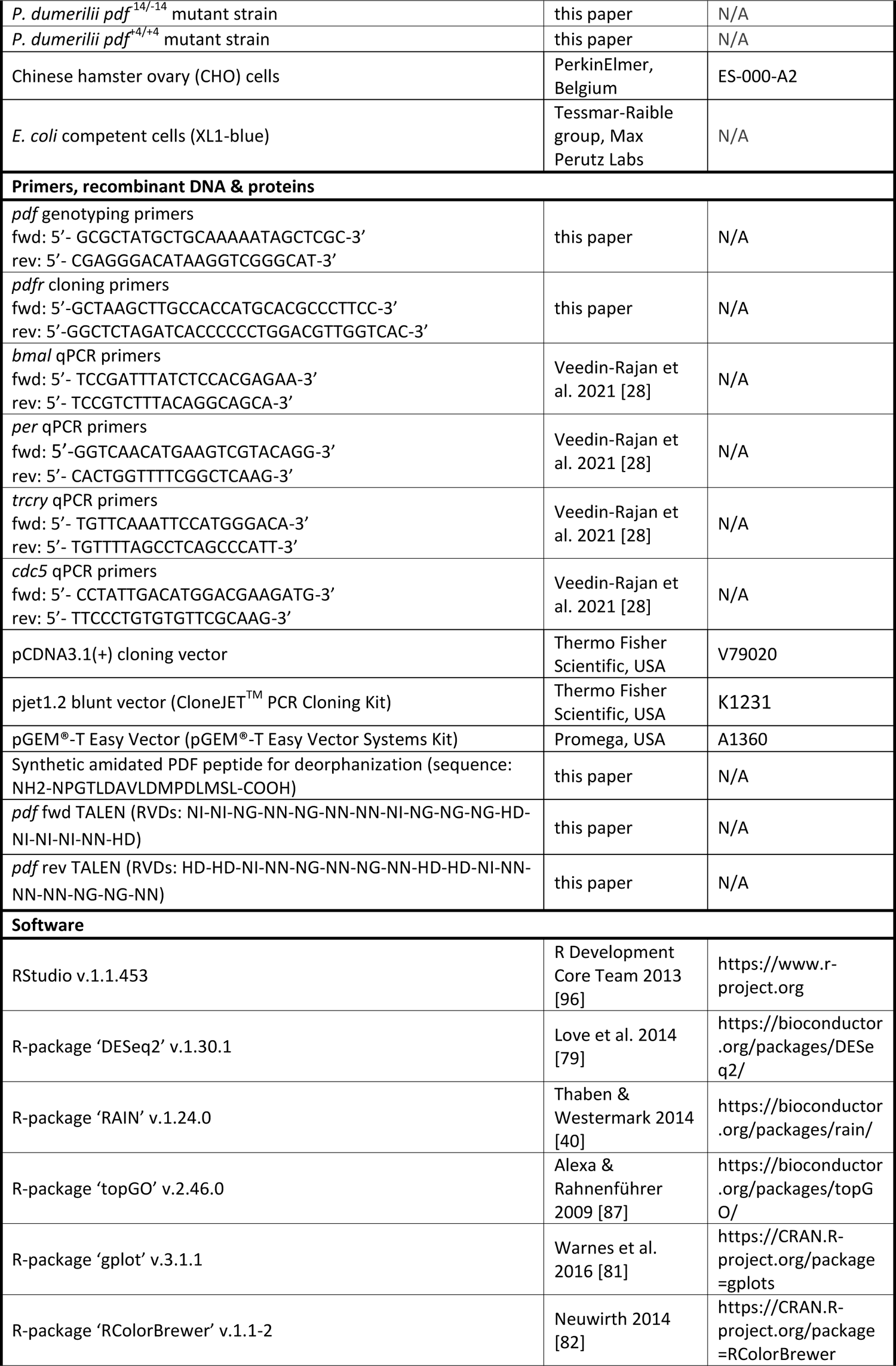
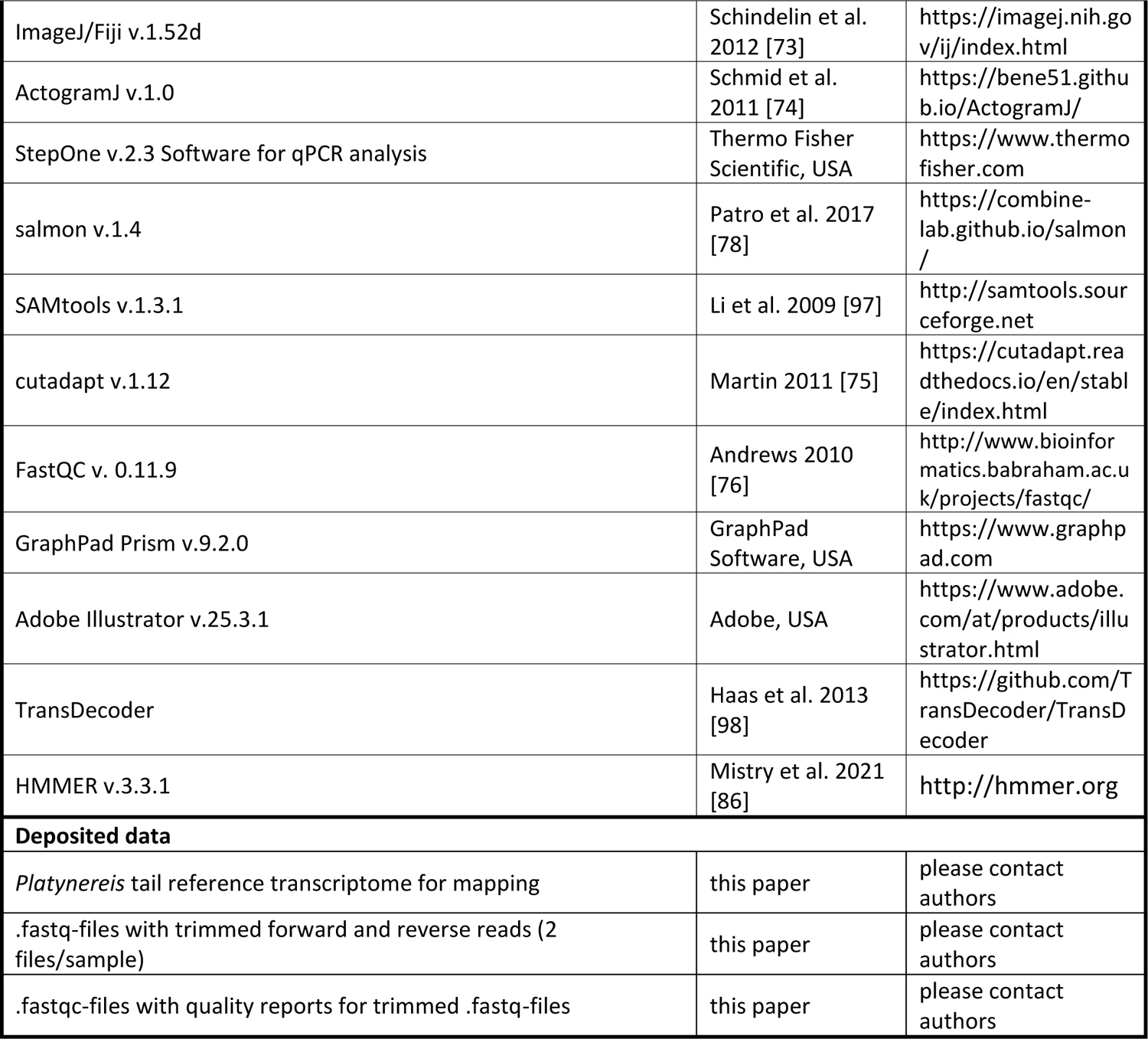

## DATA AVAILABILITY

All data relevant to the paper are provided in the supplemental material or were submitted to the Dryad online repository. For additional information on used reagents, sequence resources and bioinformatics analyses, please contact the authors directly.

## ACKNOWLEDGMENTS

N.S.H. was supported by a Lise-Meitner fellowship by the Austrian Science Fund (FWF, M2820). K.T.-R. received funding from the European Research Council under the European Community„s Seventh Framework Programme (FP7/2007–2013) ERC Grant Agreement #337011, the Horizon 2020 Programme ERC Grant Agreement #819952, and the FWF research platform „Rhythms of Life‟ of the University of Vienna (SFB F78). I.B. and L.S. received support from the KU Leuven Research Council (C16/19/003). A.M.M. was supported by the VBC post-doctoral program VIP^2^ co-funded by the Horizon 2020 Marie Skłodowska-Curie Grant Agreement #847548. F.R. acknowledges support by the FWF (#I2972), and the European Research Council (#260564). None of the funding bodies was involved in the design of the study, the collection, analysis, and interpretation of data or in writing the manuscript. We are further grateful to Andrij Belokurov, Margaryta Bosysova and Netsaneh Getachew (Max Perutz Labs Aquatic Facility) for sustaining worm cultures and routine genotyping support.

## AUTHOR CONTRIBUTIONS

N.S.H. and K.T.-R. designed the study and interpreted the results. N.S.H. wrote the manuscript with comments by K.T.-R. and review by the other coauthors. N.S.H., K.V. and A.M.M. performed locomotor activity recordings. S.H. identified the *pdf* gene sequence. K.V. generated *pdf* mutants and performed clock gene qPCRs. I.B., K.V. and L.S. performed deorphanization of the PDF-receptor. N.S.H performed rhythmic phenotype characterizations and prepared samples for RNASeq. N.S.H., F.R. and A.W.S. preprocessed transcriptome data and A.W.S. assembled the reference transcriptome used for mapping. L.H. performed transcript filtering and GO-term analysis, N.S.H. performed RAIN analysis of transcripts.

## SUPPORTING FIGURE CAPTIONS

S1 Fig: Spectral light conditions for worm incubations. (A) Standard worm culture light spectrum. (B) Behavior chamber light spectrum. (D) Logarithmic plotting of panels A and B as well as a natural sunlight spectrum (10 am - 4 pm local time average) recorded in the natural habitat of *Platynereis dumerilii* around Ischia, Italy at 5 m depth in November 2011 [36]. Overall irradiance (380-750 nm) was 4.78*10^14^ photons*cm^-^²*s^-1^ for the worm culture and 1.40*10^15^ photons*cm^-^²*s^-1^ for the behavior chamber.

S2 Fig: Individual worm actograms of strain comparison. Related to Fig 1. Double-plotted actograms of individual worms from the (A) PIN strain, (B) NAP strain and (C) VIO strain are shown. Locomotor activity was recorded over 4 LD days (16h:8h) and 8 DD days. #: individual worm identifier. Red shading indicates when worms crawled out of the tracking arena. Worms that were excluded from statistics due to maturation/spawning during or within one week after the recording are not shown, as maturation strongly alters their overall behavior. Sexes were not determined systematically, but are indicated, if known.

S3 Fig: Additional actograms of worm locomotor activity in initial and repeated runs. Related to Fig 1E-G. Double-plotted actograms of (A) PIN strain, (B) NAP strain and (C) VIO strain worms are shown. Locomotor behavior was recorded over 4 LD days (16h:8h) and 8 DD days in 2 consecutive runs (initial/repeated). # indicates the individual worm number. Red shading indicates when worms crawled out of the tracking arena. Worms that were excluded from statistics due to maturation during or within one week after the recording are not shown, as maturation strongly alters their overall behavior.

S4 Fig: Individual worm actograms of PIN wildtype characterization of RNASeq analysis. Related to Fig 2. Double-plotted actograms of individual PIN wildtype worms are shown. Locomotor activity was recorded over 3 LD days (16h:8h) and 3 DD days. Per behavioral recording, 25 worms were investigated in parallel. Two identical behavior chambers were used for recordings. Each page contains all worms of a characterization run including worms that matured/spawned and were thus excluded. #: individual worm identifier. Letters indicate characterization as rhythmic (R), arrhythmic (A) or intermediate (i).

S5 Fig: Variance of transcript subgroups with RAIN-significant cycling. Related to Fig 3. (A) Transcripts with RAIN-significant 24h cycling only in behaviorally rhythmic worms. The used transcripts (n=26) were associated with the GO-terms „neuromuscular process controlling balance‟, „axon regeneration‟, „visual behavior‟, and „response to hypoxia‟. (B) Transcripts with RAIN-significant 24h cycling only in behaviorally arrhythmic worms. The used transcripts (n=25) were associated with the GO-terms „fatty acid beta-oxidation using acyl-CoA dehydrogenase‟, „glucose metabolic process‟, „excretion‟, „response to vitamin A‟, „mitochondrial transmembrane transport‟, and „phosphatidylinositol phosphorylation‟. Mean transcript SDs for a given transcript and phenotype were calculated as mean of the SDs for the 6 individual time points (n=3 samples per time point). Variance was compared between rhythmic (blue) and arrhythmic (red) phenotypes via paired 2-sided t-test. Black lines indicated value-pairs belonging to the same transcript. Significance levels: \**p*<0.05, \*\**p*<0.01, \*\*\**p*<0.001, \*\*\*\**p*<0.0001. While there was a general trend of lower variance in the phenotype with RAIN-significant cycling, in both (A) and (B) there are transcripts with no or the opposite trend. The higher variances for neuronal/behavioral transcripts (A) in behaviorally arrhythmic worms are actually fully consistent with the observed behavior.

S6 Fig: Diel expression patterns of RAIN-significant *Platynereis dumerilii* transcripts encoding putative matrix metalloproteinases (MMPs). Related to Fig 3. Transcripts with significant 24h cycling in (A) behaviorally rhythmic worms, (B) behaviorally arrhythmic worms, or (C) both are shown for both phenotypes, respectively (rhythmic: blue, arrhythmic: red). Bold *p*-values indicate significant 24h cycling as determined by RAIN analysis. Per time point, n=3 replicates were measured. Further details on MMP transcripts are provided in S3 Tab.

S7 Fig: Individual worm actograms of *pdf* wildtype/mutant comparison. Shown are double-plotted actograms of mixed VIO/PIN background *pdf* wildtypes (A) and mutants (B) related to Fig 6, the initial VIO background wildtypes (C) and mutants (D) related to S8 Fig, and VIO backcrossed wildtypes (E) and mutants (F) related to S9 Fig. Locomotor activity was recorded over 4 LD days (16h:8h) and 4,5 or 8 DD days. #: individual worm identifier. Genotypes of *pdf* mutants (−14/-14, +4/+4, −14/+4) are indicated at the bottom of the respective actograms. Red shading indicates when worms crawled out of the tracking arena. Worms that were excluded from statistics due to maturation/spawning during or within one week after the recording are not shown, as maturation strongly alters their overall behavior.

S8 Fig: behavioral rhythmicity of *pdf* wildtypes & mutant in the initial VIO strain background. Related to Fig 6. In four recordings, n=34 *pdf* wildtypes from 4 mating batches were compared to n=43 *pdf* mutants from 5 mating batches (−14/-14 n=33, −14/+4 n=10). (A) Circadian locomotor activity of VIO strain *pdf* wildtypes (black) and mutants (red) under 4 days of LD and 8 days of DD. Individual worm actograms are provided in S7C,D Fig. (B) Cumulative activity over the early day (0-8), late day (8-16) and night (16-24) in LD and DD. (C,D) Period/power of wildtype and mutant locomotor rhythms in the circadian range (20h-28h) in LD and DD determined by Lomb-Scargle periodogram. Statistical differences (panel B-D) were determined via Mann-Whitney U-test. For further info on figure labeling see main text Fig 1. The reduced rhythmicity of *pdf* mutants disappeared directly after an outcross against the PIN strain (Fig 6) and could not be recovered by backcross against the VIO strain (S9 Fig). Hence we consider it to be an off-target effect, i.e. not causally connected to the *pdf* mutant locus.

S9 Fig: behavioral rhythmicity of *pdf* wildtypes & mutant after backcross into the VIO strain. Related to Fig 6. In one recording, n=19 *pdf* wildtypes from 5 mating batches were compared to n=19 *pdf* mutants from 6 mating batches (−14/-14 n=9, +4/+4 n=10). (A) Circadian locomotor activity of VIO-backcrossed *pdf* wildtypes (black) and mutants (red) under 4 days of LD and 8 days of DD. Individual worm actograms were are provided in S7E,F Fig). (B) Cumulative activity over the early day (0-8), late day (8-16) and night (16-24) in LD and DD. (C,D) Period/power of wildtype and mutant locomotor rhythms in the circadian range (20h-28h) in LD and DD determined by Lomb-Scargle periodogram. Statistical differences were determined via Mann-Whitney U-test (panel B,C) or unpaired 2-sided t-test (panel D). For further info on figure labeling see main text Fig 1. The persistence of the phenotype observed in PIN-outcrossed worms (Fig 6) reinforces that this is a solid *pdf* mutant phenotype and that the initial rhythmicity reduction in the VIO strain (S8 Fig) is not causally connected to the *pdf* mutant locus.

