## Supplemental Text 1 for "Behavioral rhythms are bad predictors of general organismal rhythmicity"

S1 Text: Details on transcript oscillation amplitudes in behaviorally rhythmic versus arrhythmic *Platynereis* wildtype worms.

For the heatmaps (Fig 3B-D), counts of a given transcript were normalized together across rhythmic and arrhythmic worms in order to allow for a direct comparison of this transcript between the different phenotypes. This plotting of the RAIN-significant transcripts revealed that transcripts significantly cycling in either rhythmic (Fig 3B) or arrhythmic worms (Fig 3D) sometimes, but not always showed similar cyclic tendencies in the respective other phenotype (Fig 3B-D). As illustrated by the heatmaps, the distinction by RAIN between cyclic and non-cyclic expression was in many cases related to oscillation amplitudes. These differences in amplitude could in principle arise in two ways. Scenario 1: Individual worms could still be rhythmic for a given transcript over time, but the phases of this transcript would be less synchronized across different worms. As we sampled multiple worm heads for each time point, this would – due to averaging – result in a seemingly lowered amplitude or even the loss of rhythmicity for a given transcript. Scenario 2: A given transcript could indeed oscillate at lower amplitude or exhibit no diel oscillations in the respective phenotype.

The two scenarios should be discernable based on their data variance. A reduction of phase synchronization of transcript between different worms while maintaining amplitude (Scenario 1) should result in an overall increased variance between biological replicates (i.e. standard deviation). If the transcripts indeed cycle less in individual worms (Scenario 2), the variance should overall not be affected across the different biological replicates. The inspection of subgroups of transcripts significantly cycling only in rhythmic worms (GO-terms related to behavior or neuronal processes) or only in arrhythmic worms (GO-terms related to metabolism) showed that indeed both scenarios can be found (S5 Fig).
