## Supplementary figures and images for "Behavioral rhythms are bad predictors of general organismal rhythmicity"

### Supplemental Figure 1

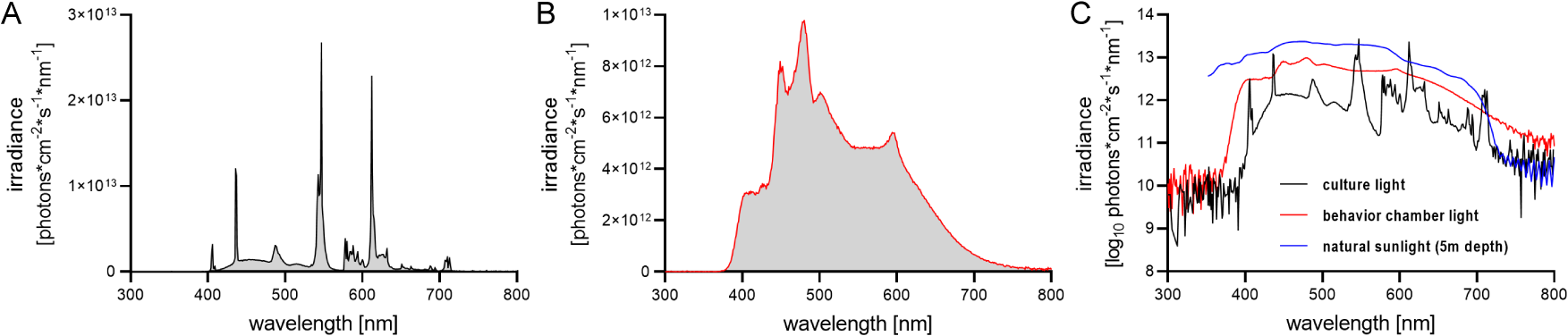

### Supplemental Figure 3

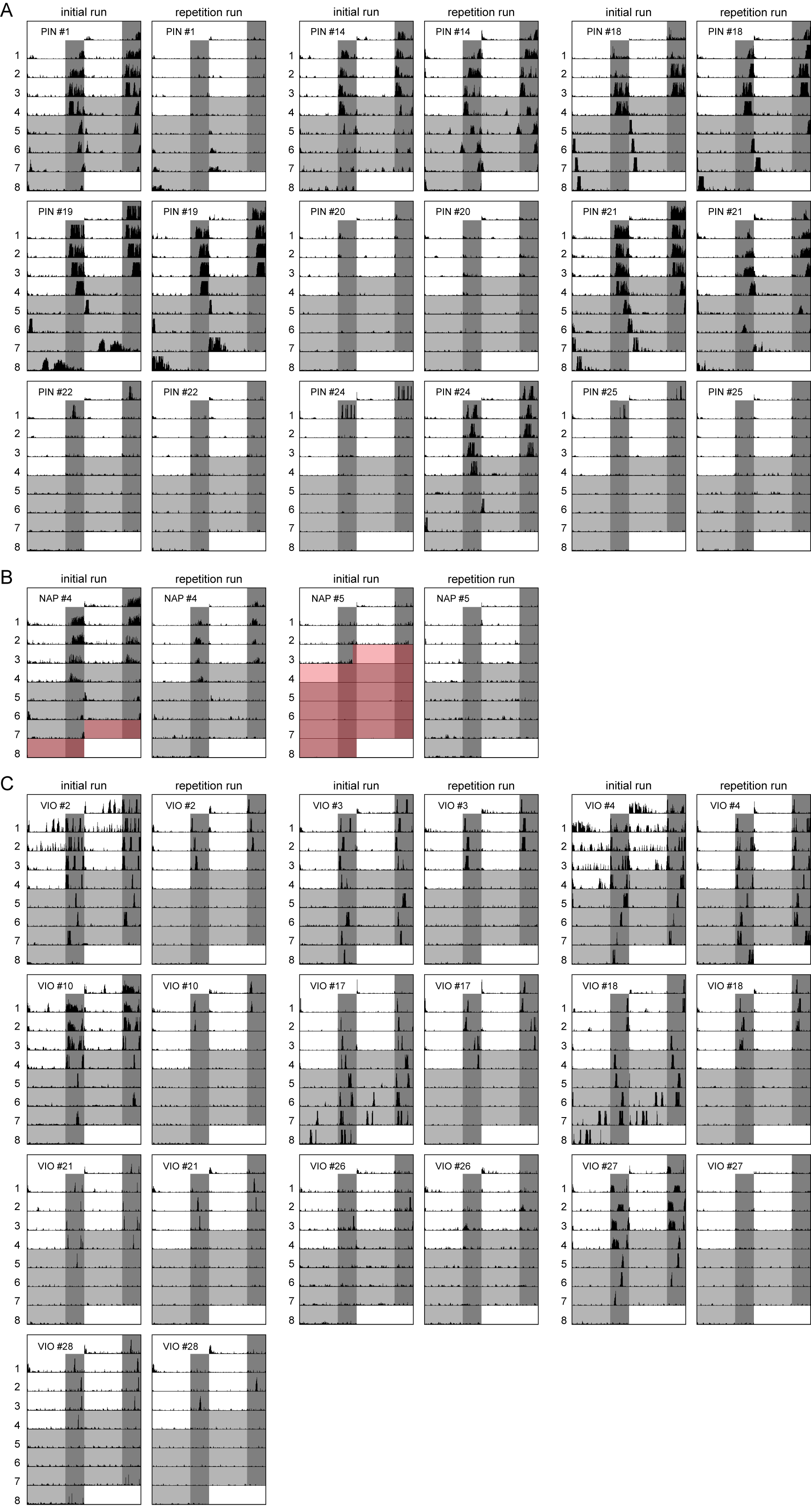

### Supplemental Figure 5

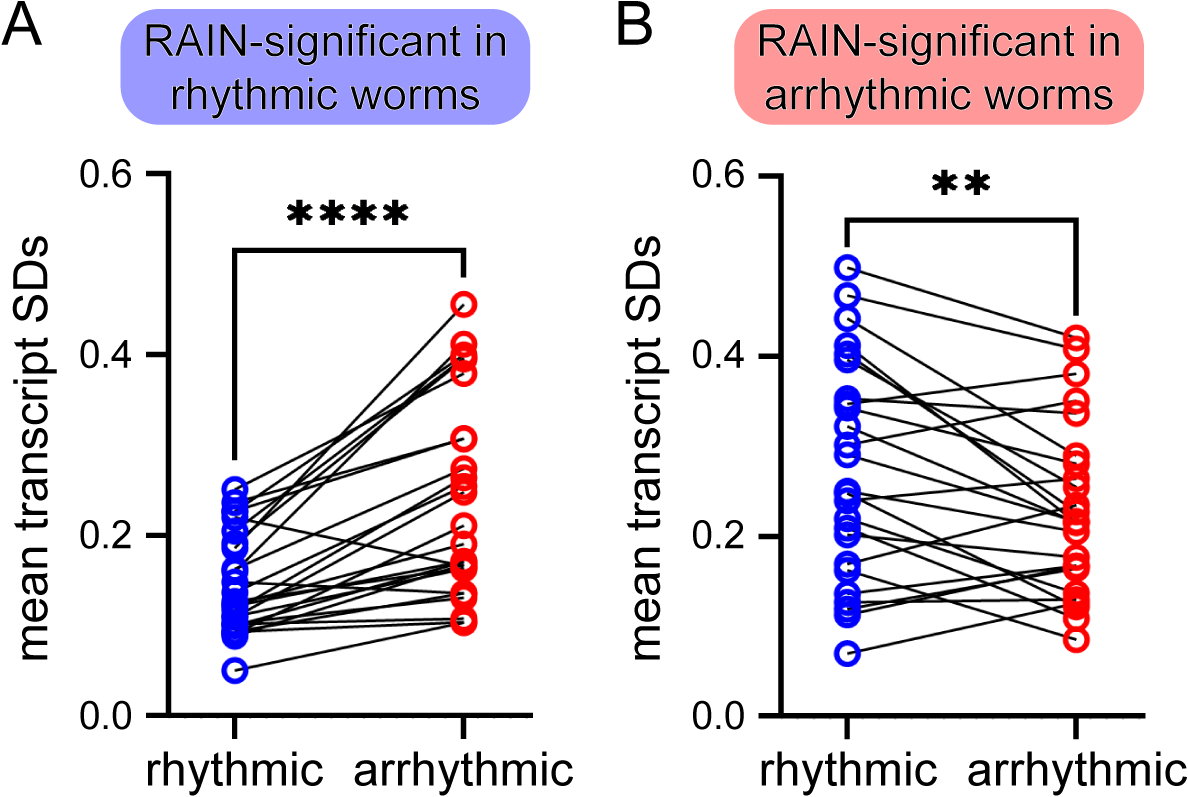

### Supplemental Figure 6

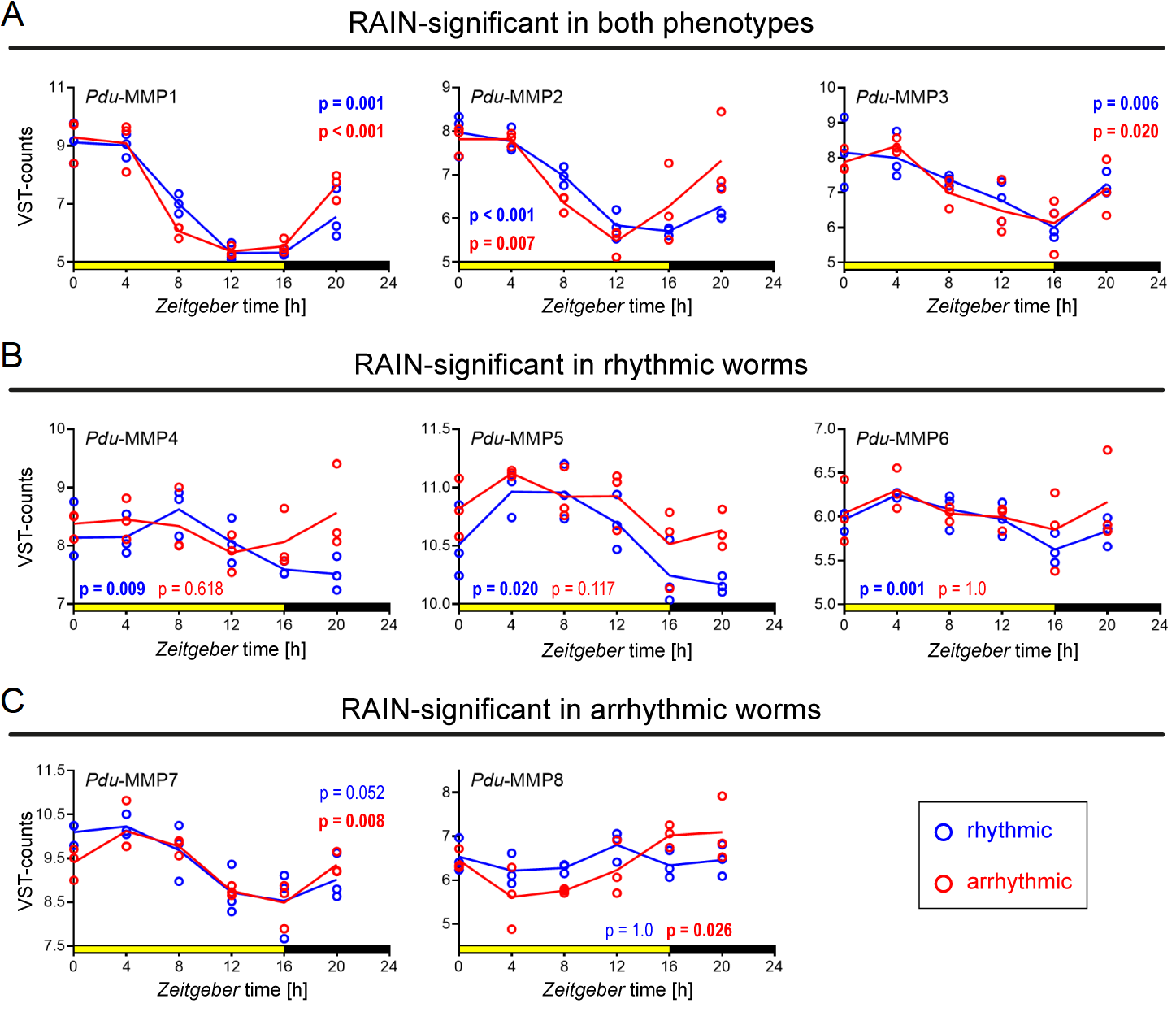

### Supplemental Figure 8

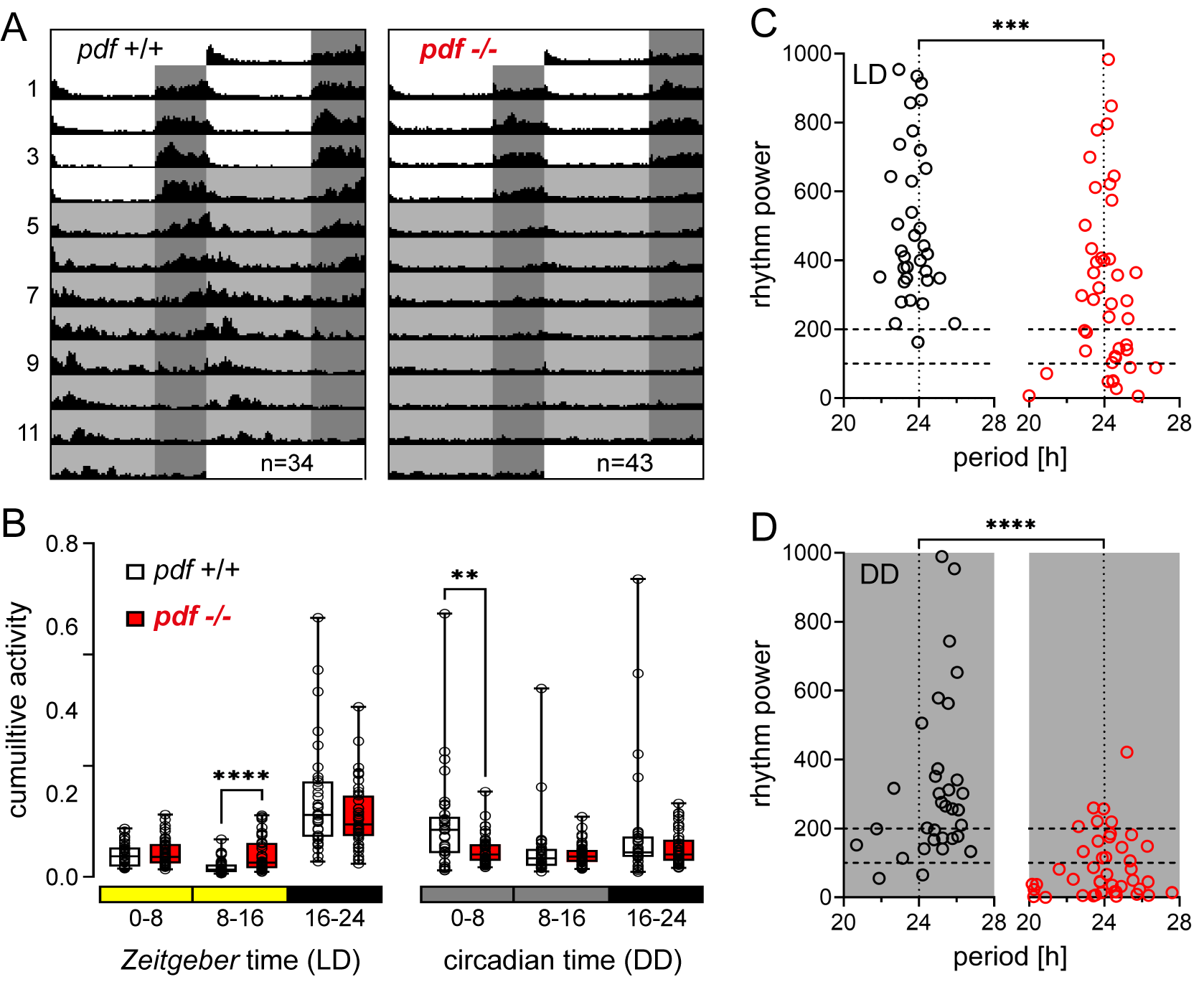

### Supplemental Figure 9

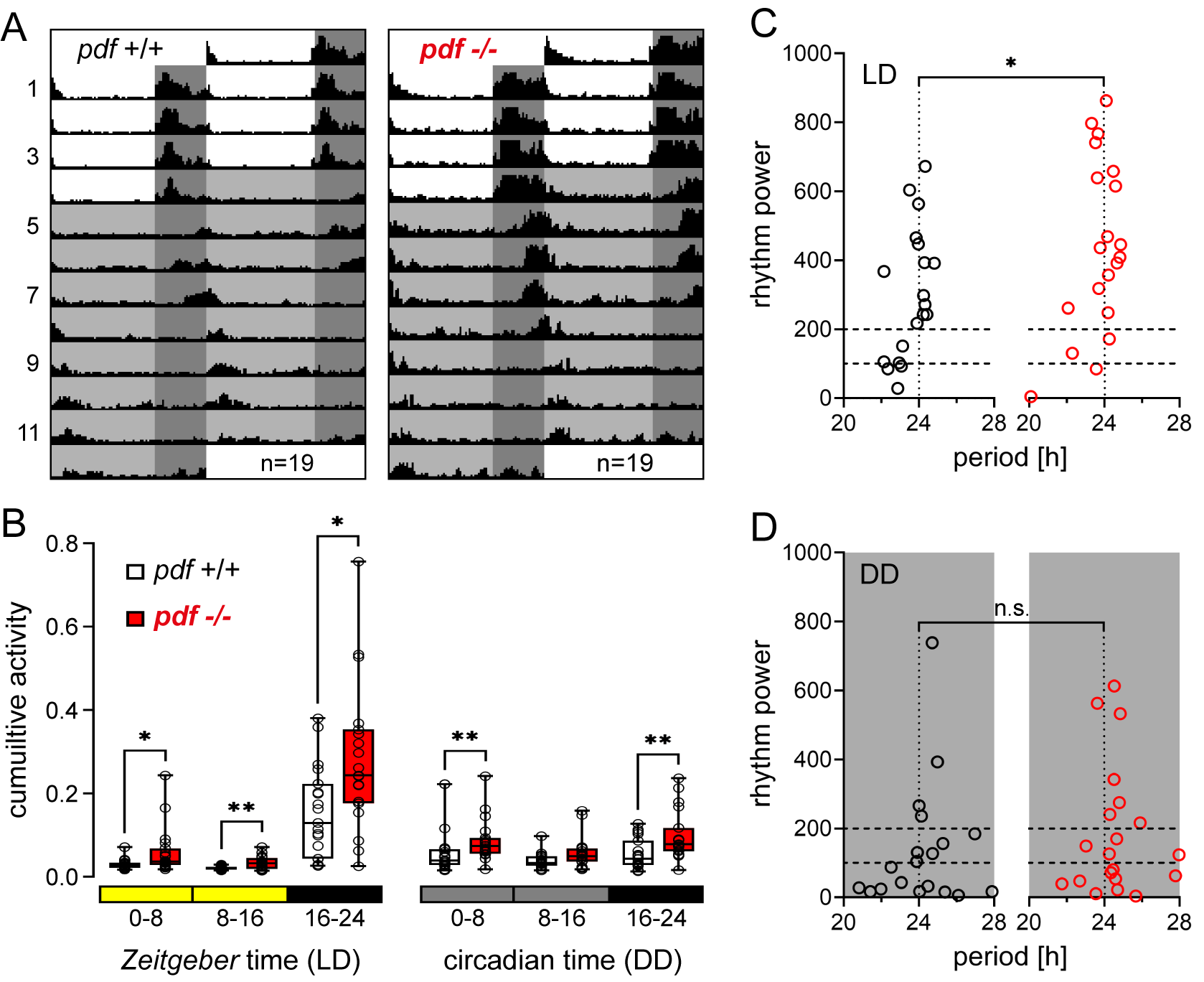
