## Supplemental Figure 2 for "Behavioral rhythms are bad predictors of general organismal rhythmicity"

<u>S2 Fig: Individual worm actograms of strain comparison. Related to Fig 1.</u> Double-plotted actograms of individual worms from the (A) PIN strain, (B) NAP strain and (C) VIO strain are shown. Locomotor activity was recorded over 4 LD days (16h:8h) and 8 DD days. #: individual worm identifier. Red shading indicates when worms crawled out of the tracking arena. Worms that were excluded from statistics due to maturation/spawning during or within one week after the recording are not shown, as maturation strongly alters their overall behavior. Sexes were not determined systematically, but are indicated, if known.

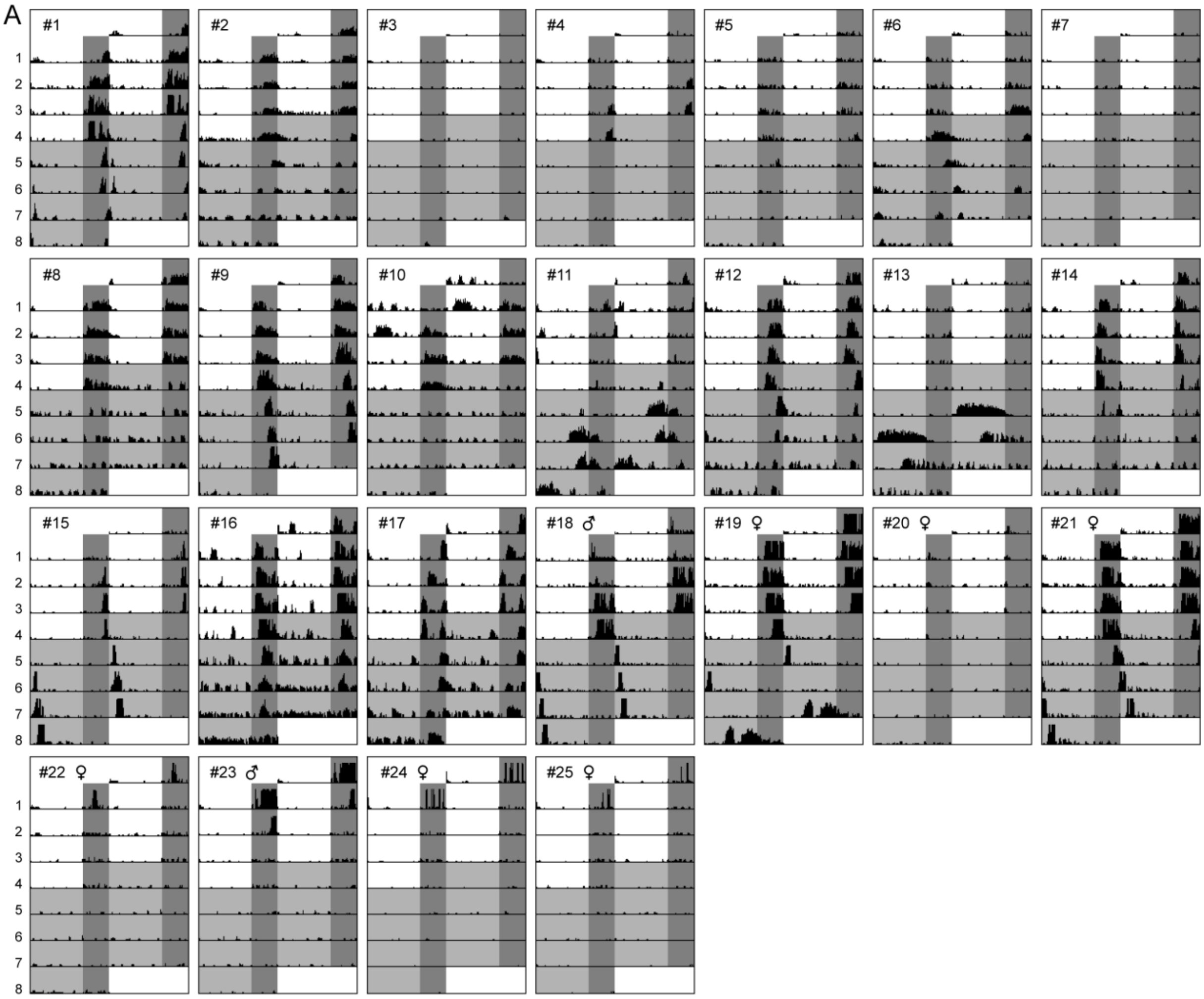

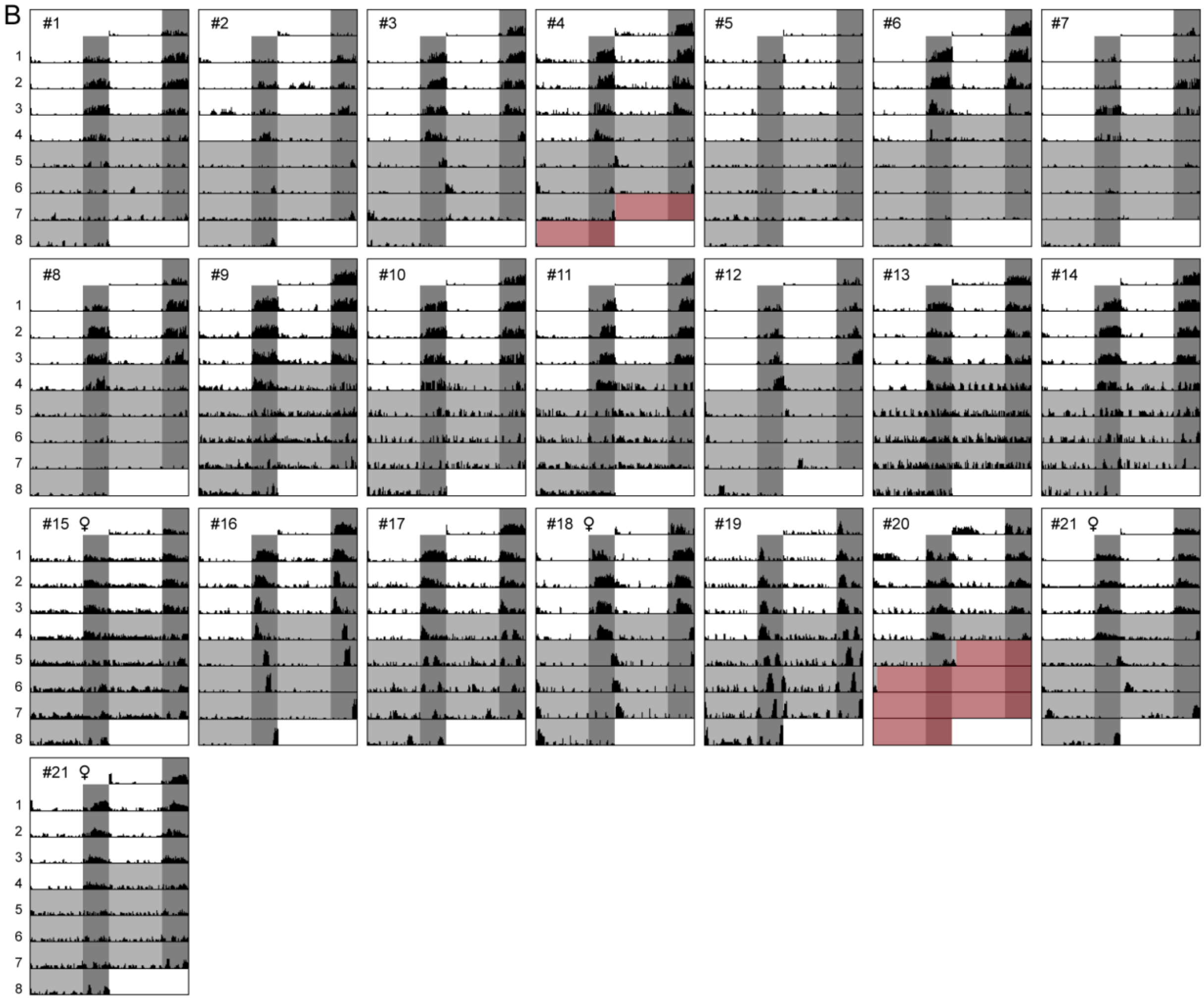

B
#1
#2
#3
#4
#5
#6
#7
#8
#9
#10
#11
#12
#13
#14
#15 ♀
#16
#17
#18 ♀
#19
#20
#21 ♀
#21 ♀
1
2
3
4
5
6
7
8

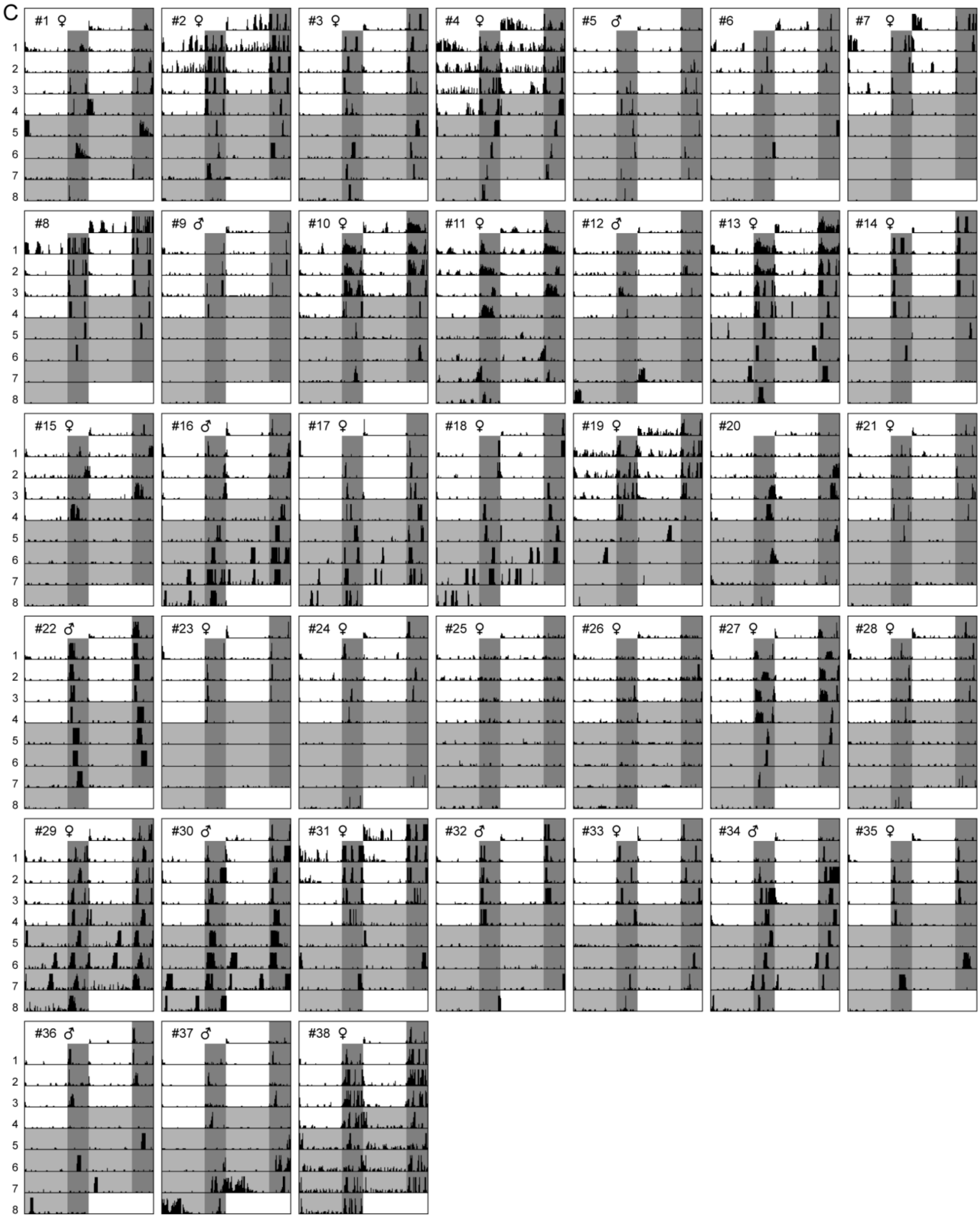
C
#1 ♀
#2 ♀
#3 ♀
#4 ♀
#5 ♂
#6
#7 ♀
#8
#9 ♂
#10 ♀
#11 ♀
#12 ♂
#13 ♀
#14 ♀
#15 ♀
#16 ♂
#17 ♀
#18 ♀
#19 ♀
#20
#21 ♀
#22 ♂
#23 ♀
#24 ♀
#25 ♀
#26 ♀
#27 ♀
#28 ♀
#29 ♀
#30 ♂
#31 ♀
#32 ♂
#33 ♀
#34 ♂
#35 ♀
#36 ♂
#37 ♂
#38 ♀
