## Supplemental Figure 4 for "Behavioral rhythms are bad predictors of general organismal rhythmicity"

<u>S4 Fig: Individual worm actograms of PIN wildtype characterization of RNASeq analysis. Related to Fig 2.</u> Double-plotted actograms of individual PIN wildtype worms are shown. Locomotor activity was recorded over 3 LD days (16h:8h) and 3 DD days. Per behavioral recording, 25 worms were investigated in parallel. Two identical behavior chambers were used for recordings. Each page contains all worms of a characterization run including worms that matured/spawned and were thus excluded. #: individual worm identifier. Letters indicate characterization as rhythmic (R), arrhythmic (A) or intermediate (i).

recording #1 (02.11.2019-07.11.2019, chamber 1)

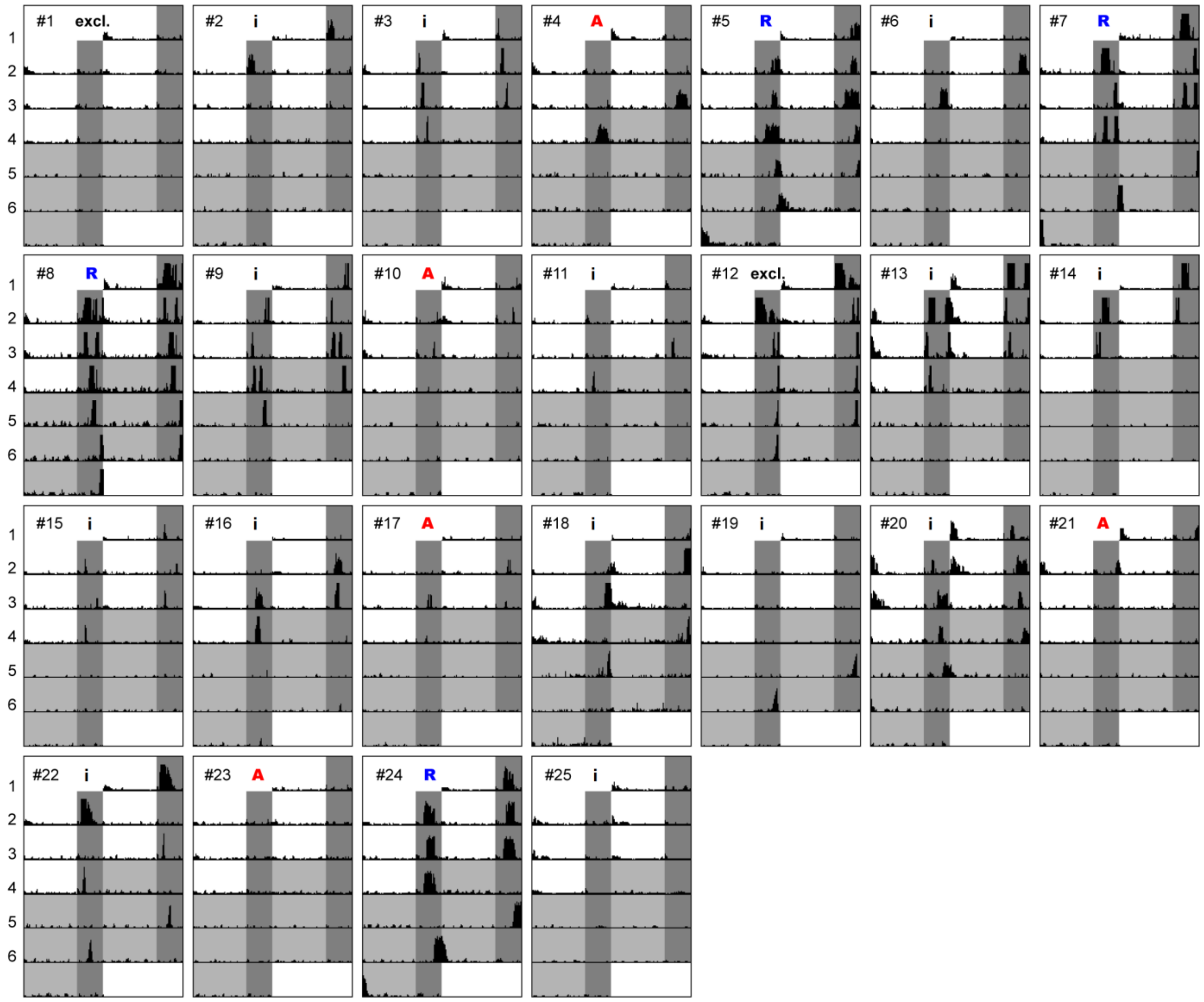

recording #2 (06.11.2019-13.11.2019, chamber 2)

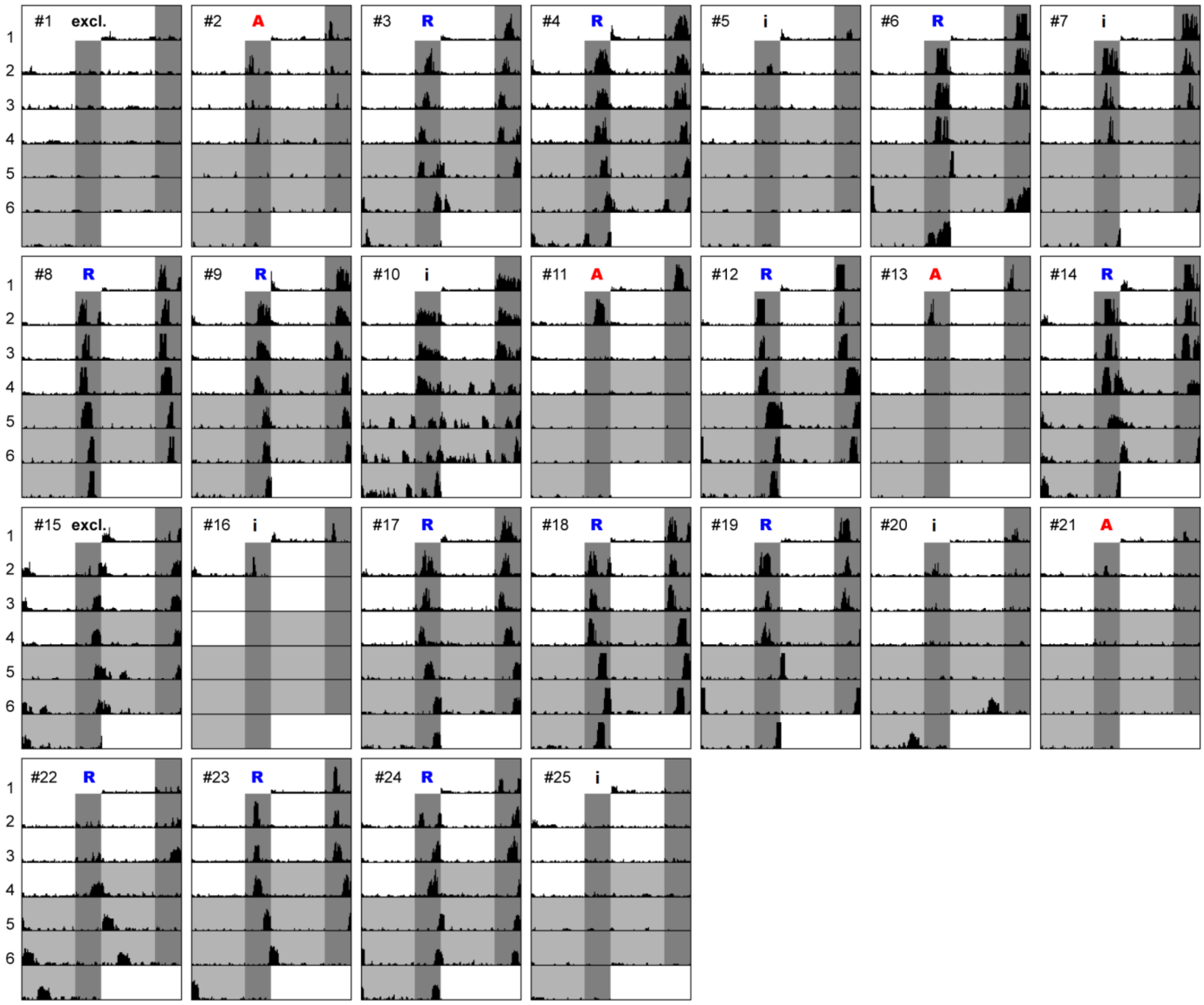

recording #3 (20.11.2019-25.11.2019, chamber 1)

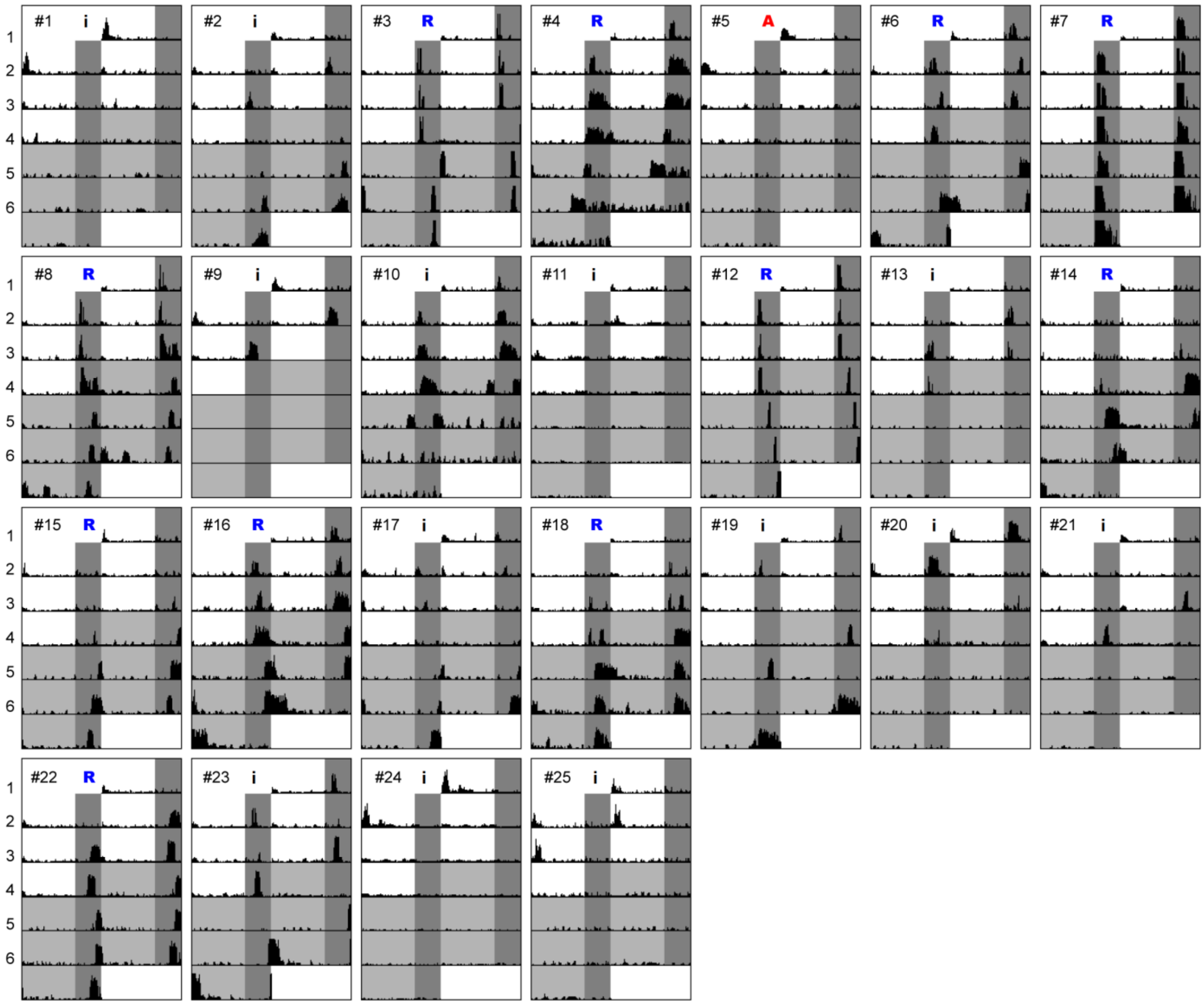

recording #4 (20.11.2019-25.11.2019, chamber 2)

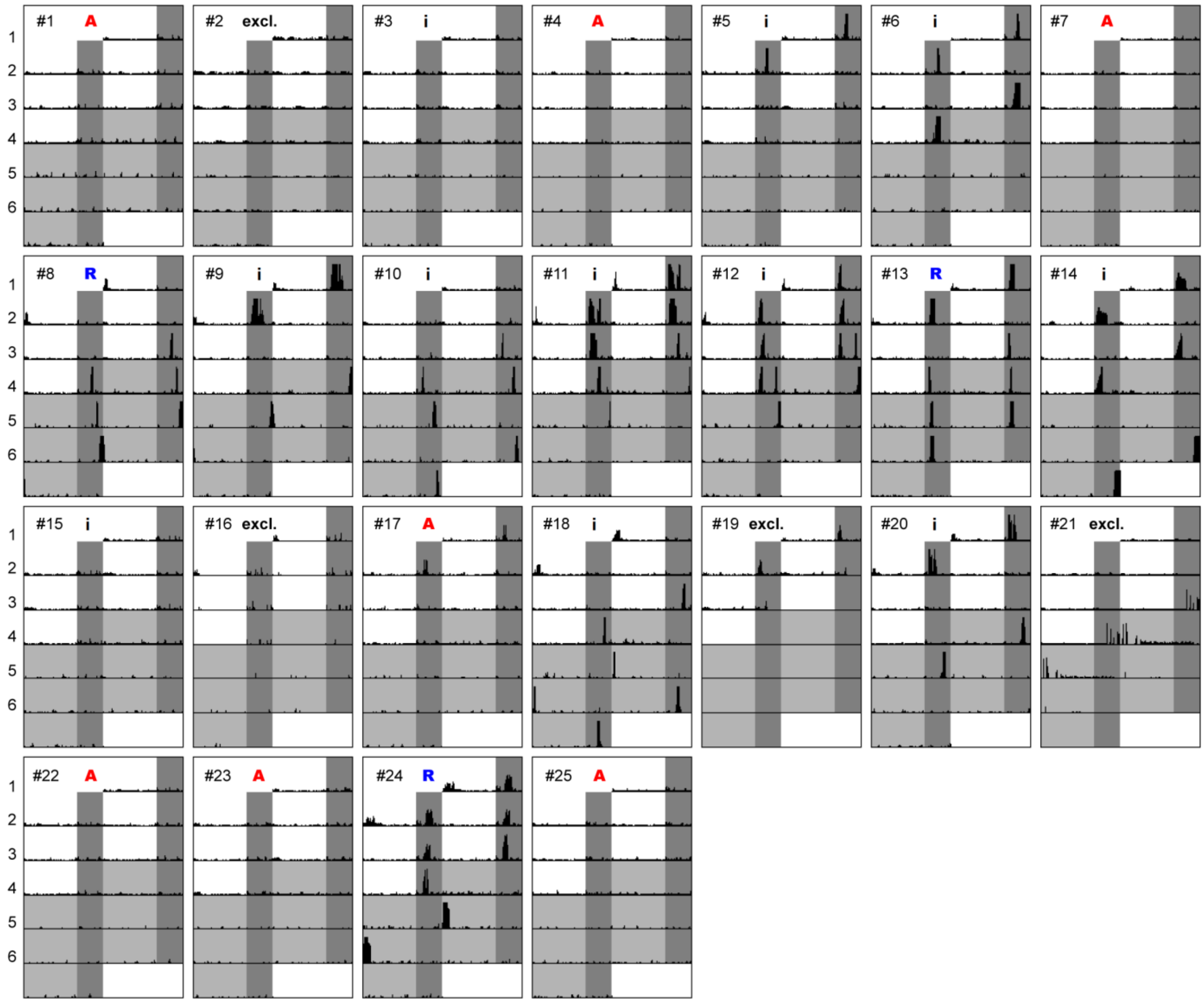

recording #5 (26.01.20-31.01.2020, chamber 1)

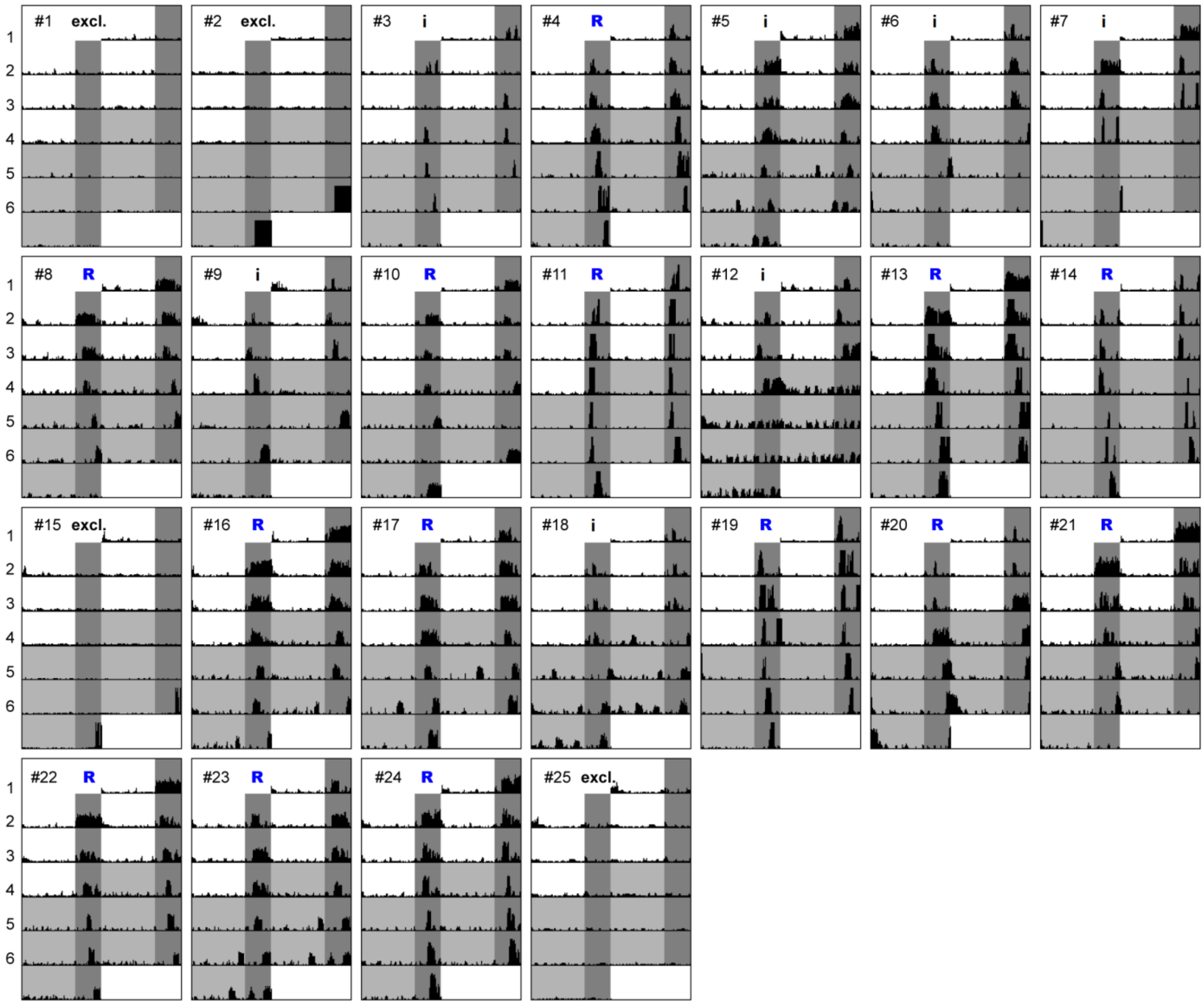

recording #6 (26.01.20-31.01.2020, chamber 2)

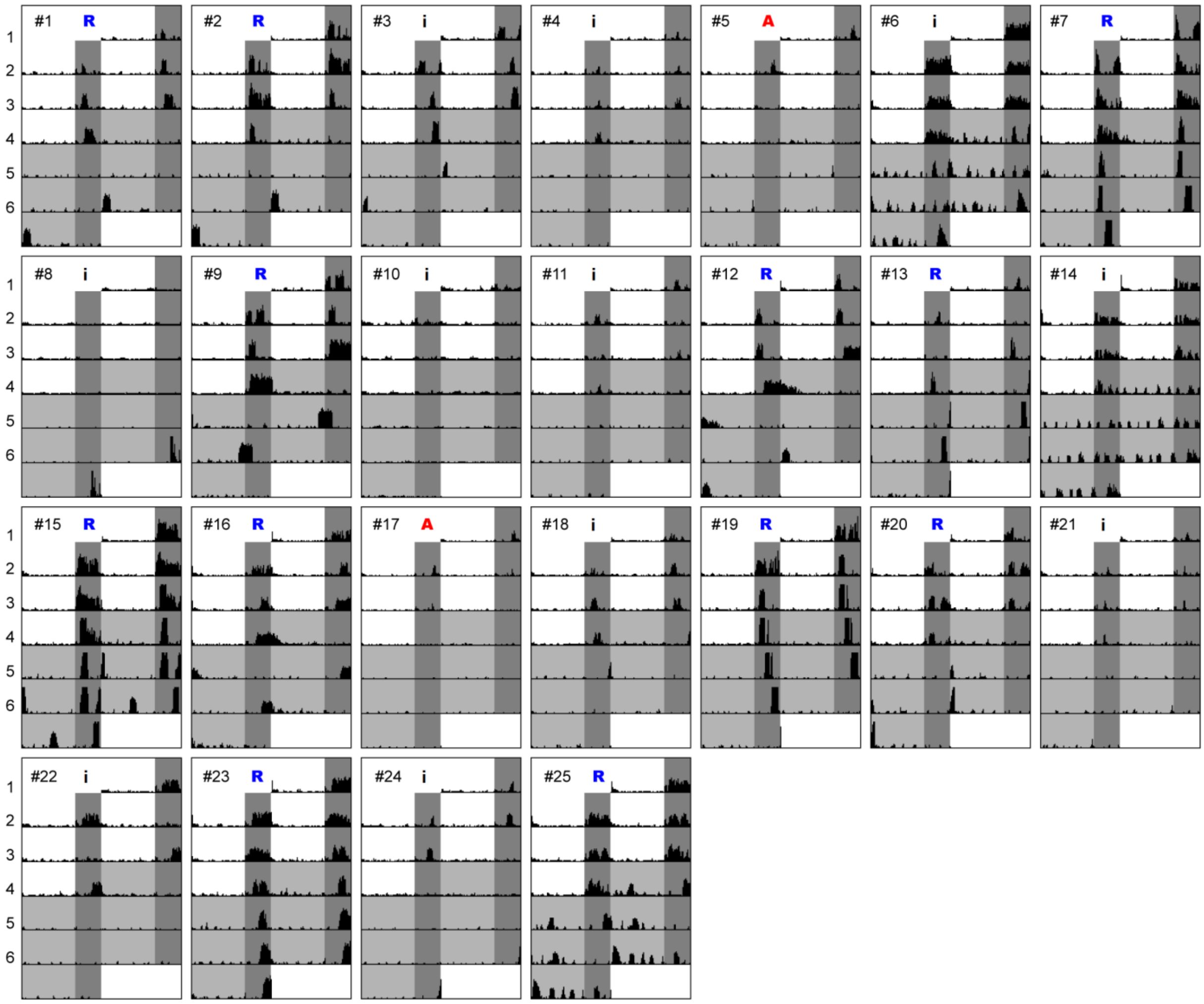

recording #7 (18.02.20-23.02.2020, chamber 1)

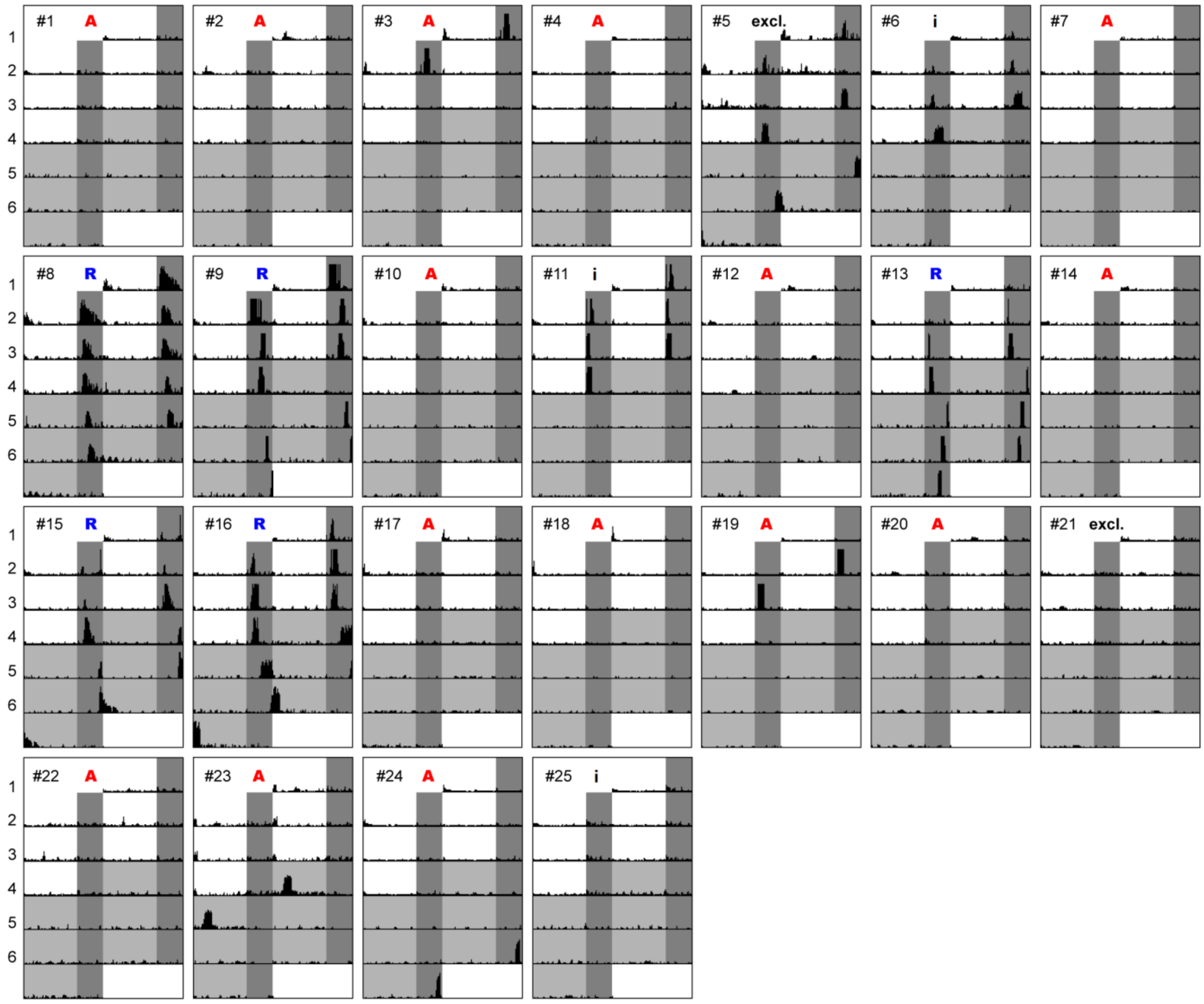

recording #8 (18.02.20-23.02.2020, chamber 2)

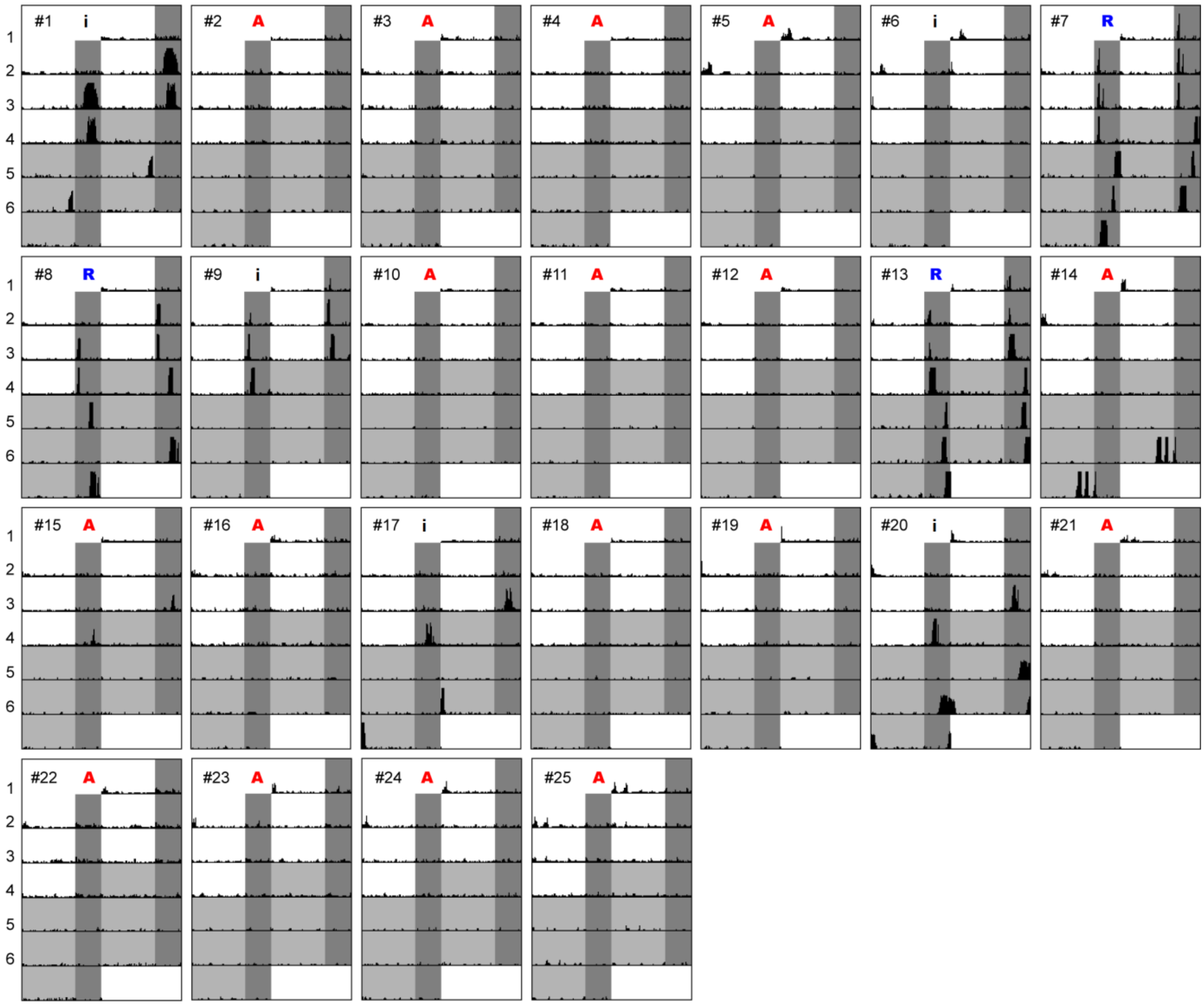

recording #9 (25.02.20-01.03.2020, chamber 1)

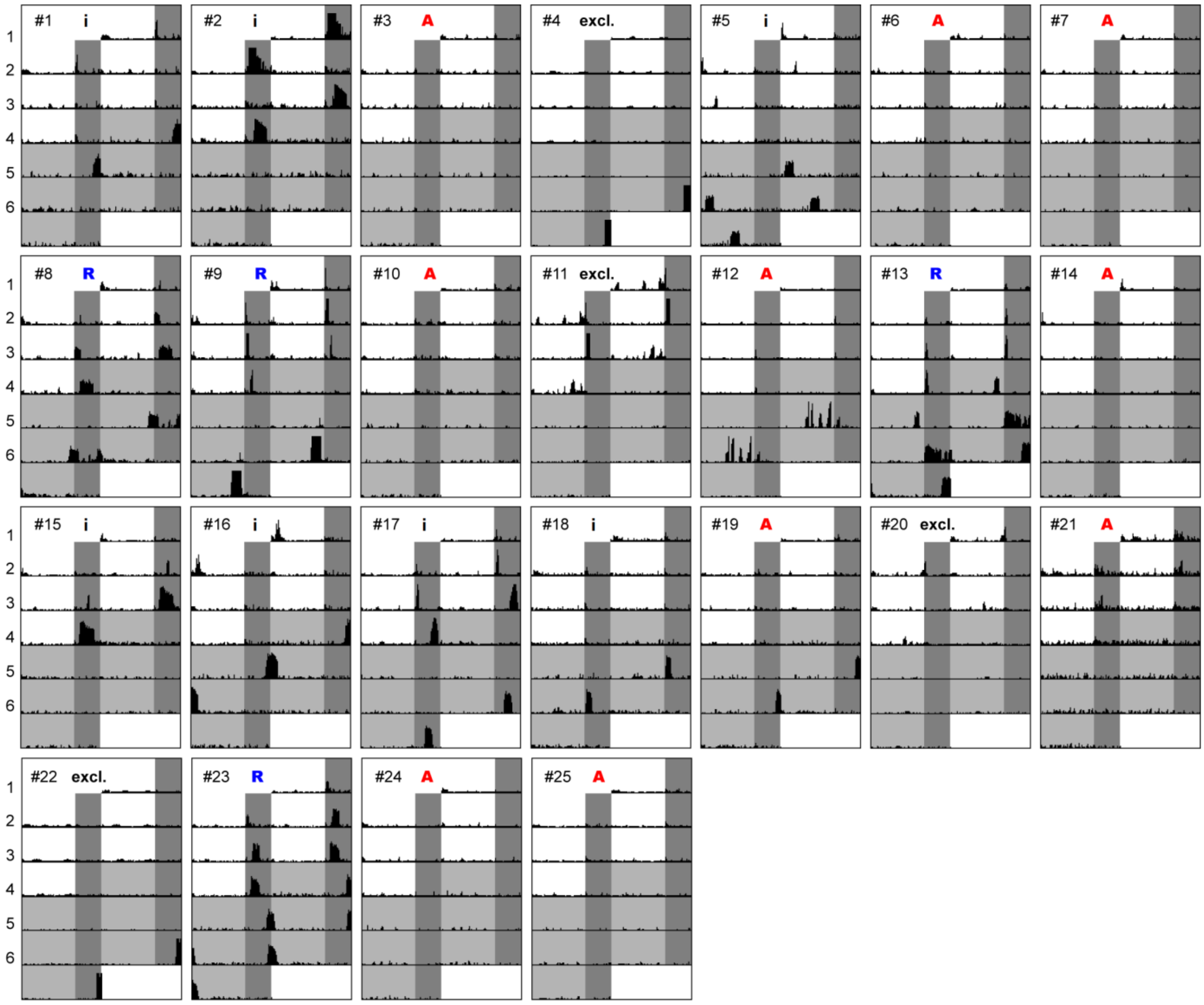

recording #10 (27.03.20-02.04.2020, chamber 1)

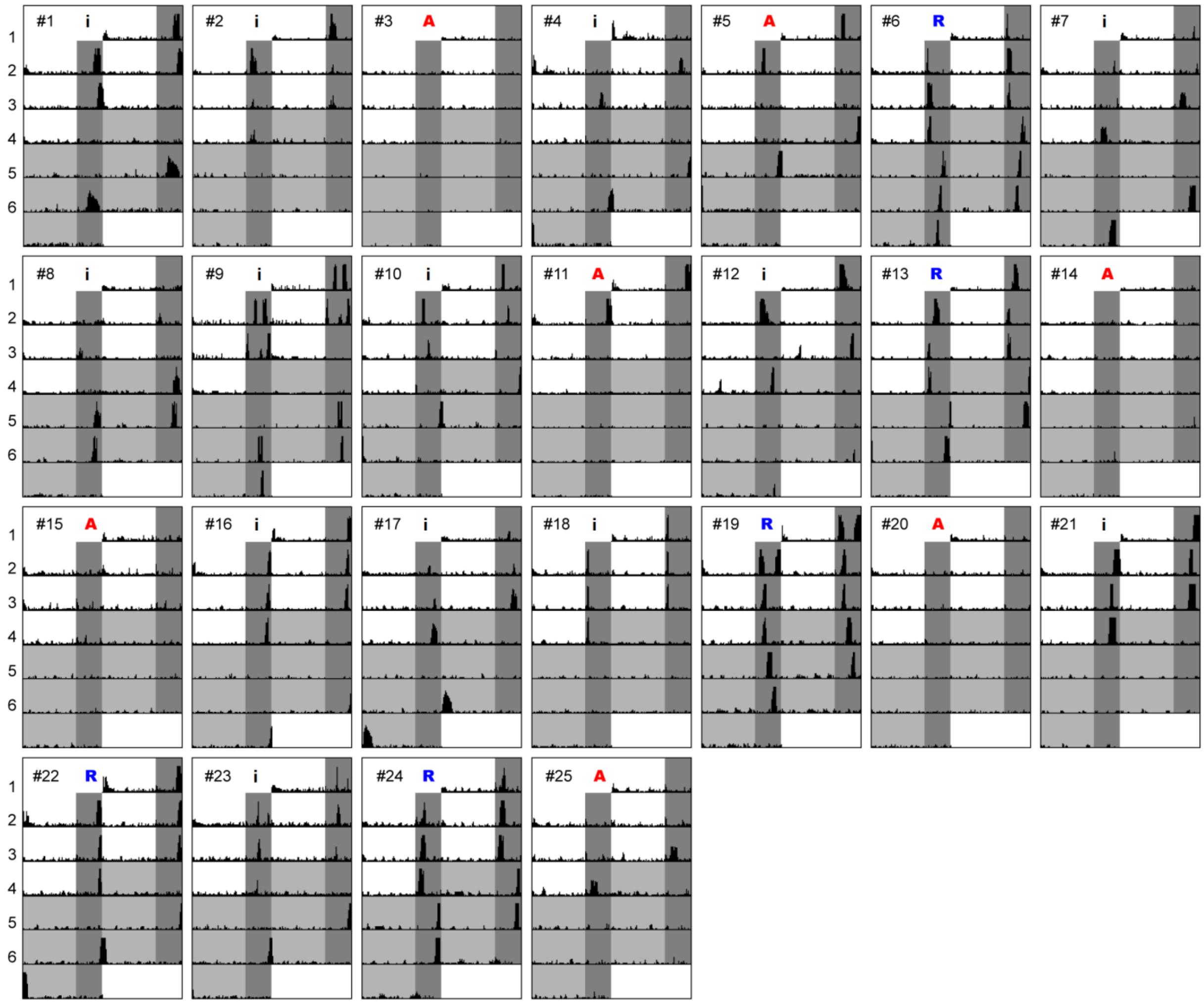

recording #11 (27.03.20-02.04.2020, chamber 2)

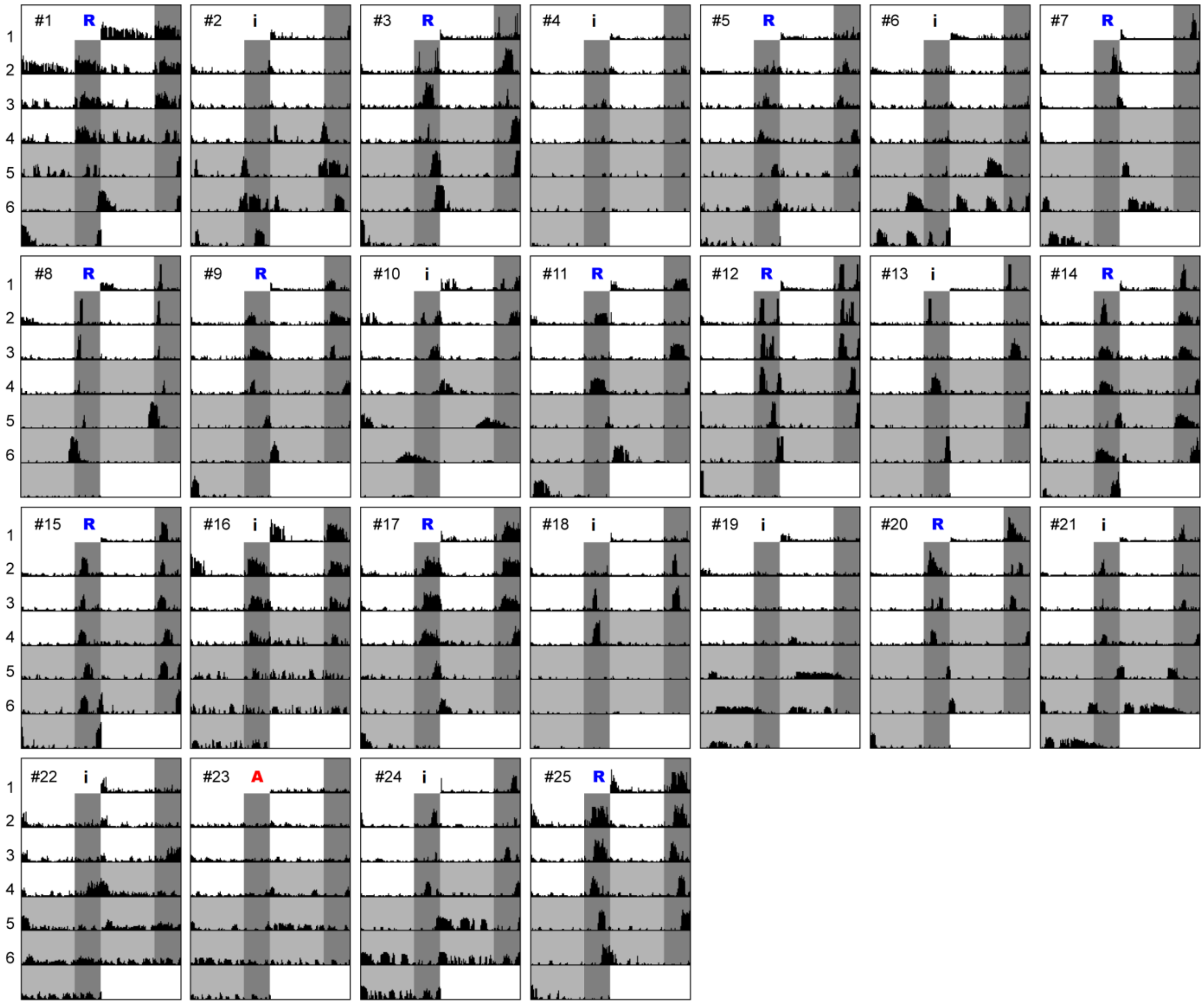

recording #12 (23.05.20-29.05.2020, chamber 1)

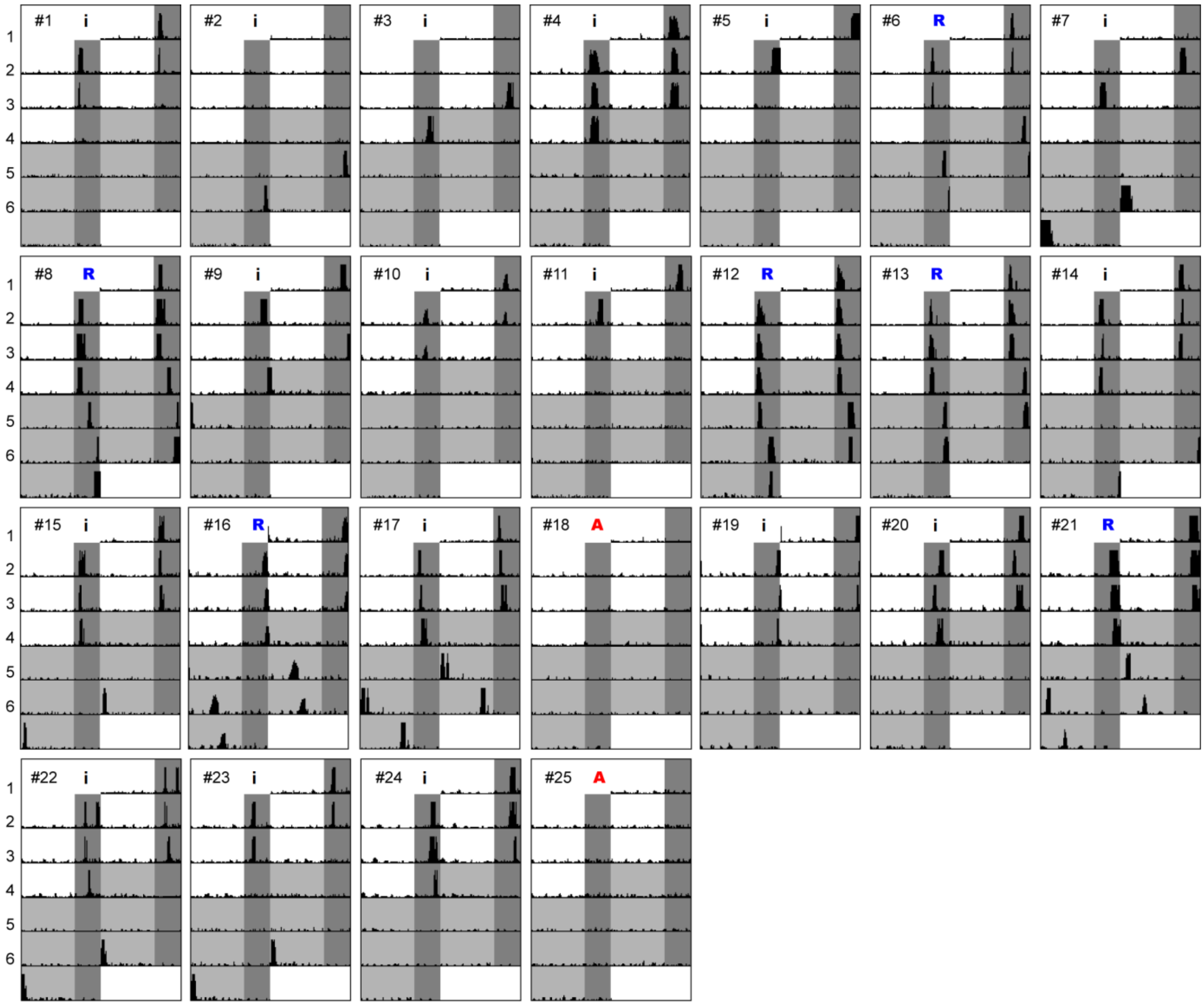

recording #13 (23.05.20-29.05.2020, chamber 2)

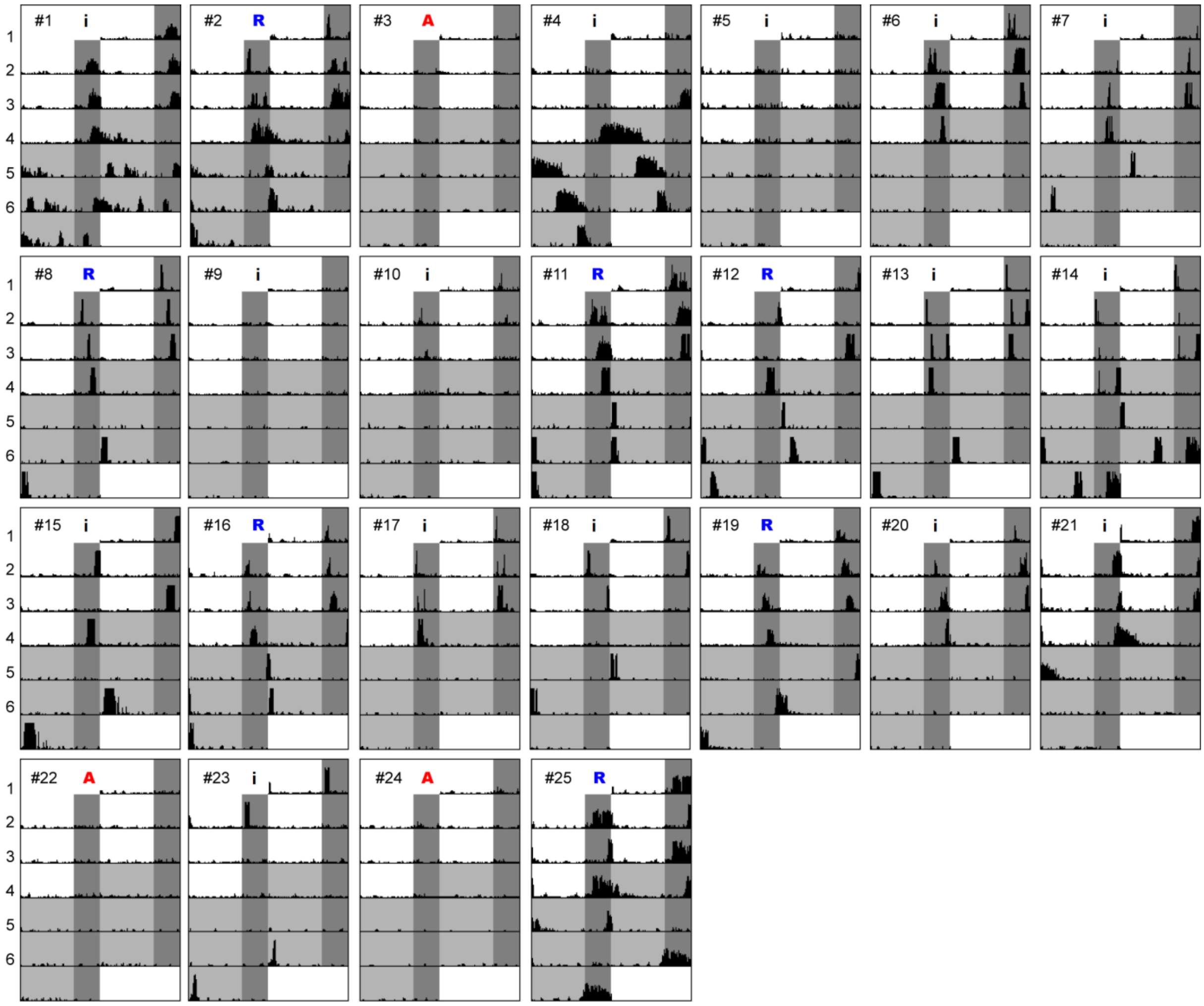
