## Supplemental Figure 7 for "Behavioral rhythms are bad predictors of general organismal rhythmicity"

<u>S7 Fig: Individual worm actograms of *pdf* wildtype/mutant comparison.</u> Shown are double-plotted actograms of mixed VIO/PIN background *pdf* wildtypes (A) and mutants (B) related to Fig 6, the initial VIO background wildtypes (C) and mutants (D) related to Fig S8, and VIO backcrossed wildtypes (E) and mutants (F) related to Fig S9. Locomotor activity was recorded over 4 LD days (16h:8h) and 4,5 or 8 DD days. #: individual worm identifier. Genotypes of *pdf* mutants (-14/-14, +4/+4, -14/+4) are indicated at the bottom of the respective actograms. Red shading indicates when worms crawled out of the tracking arena. Worms that were excluded from statistics due to maturation/spawning during or within one week after the recording are not shown, as maturation strongly alters their overall behavior.

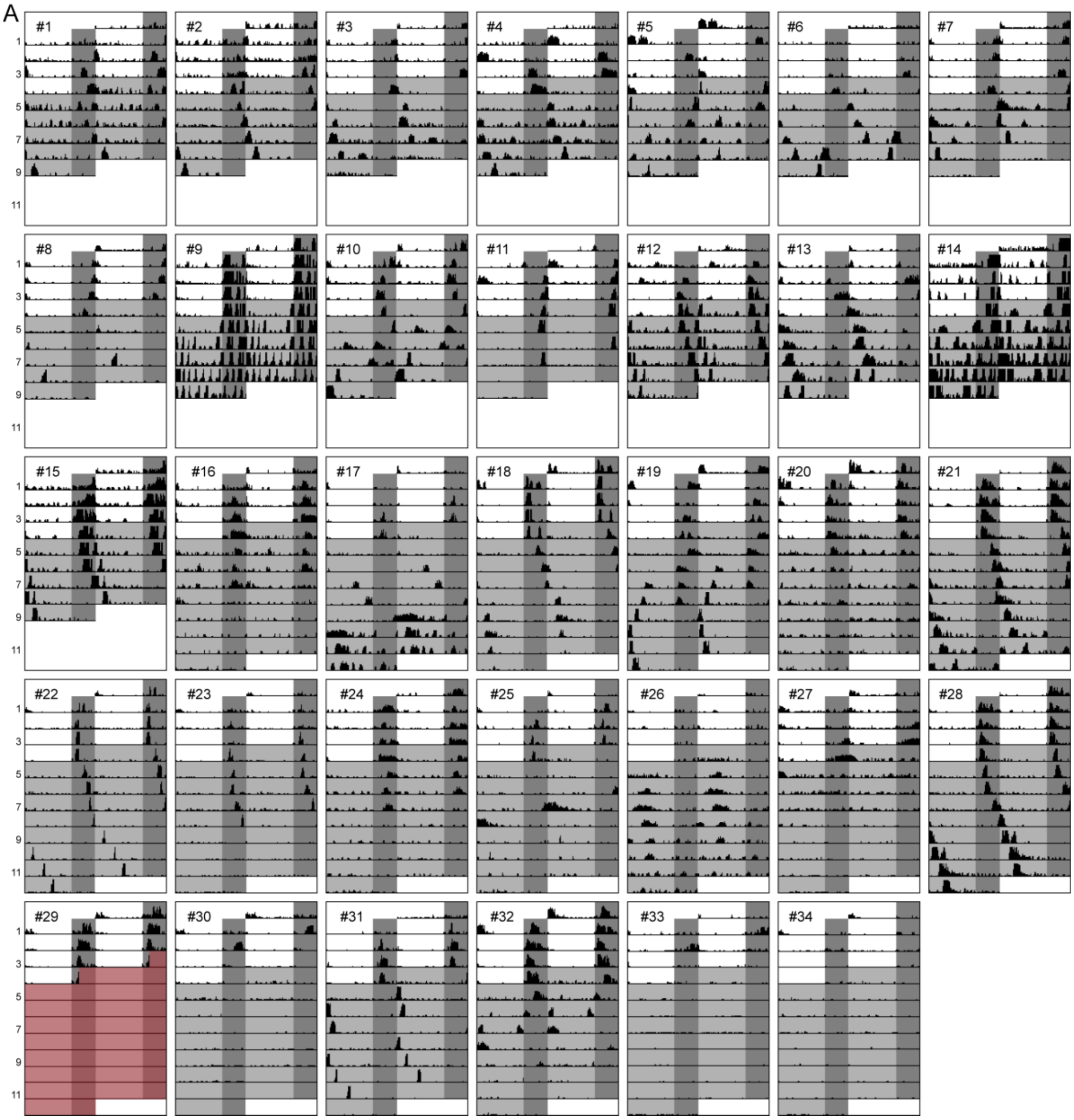

B

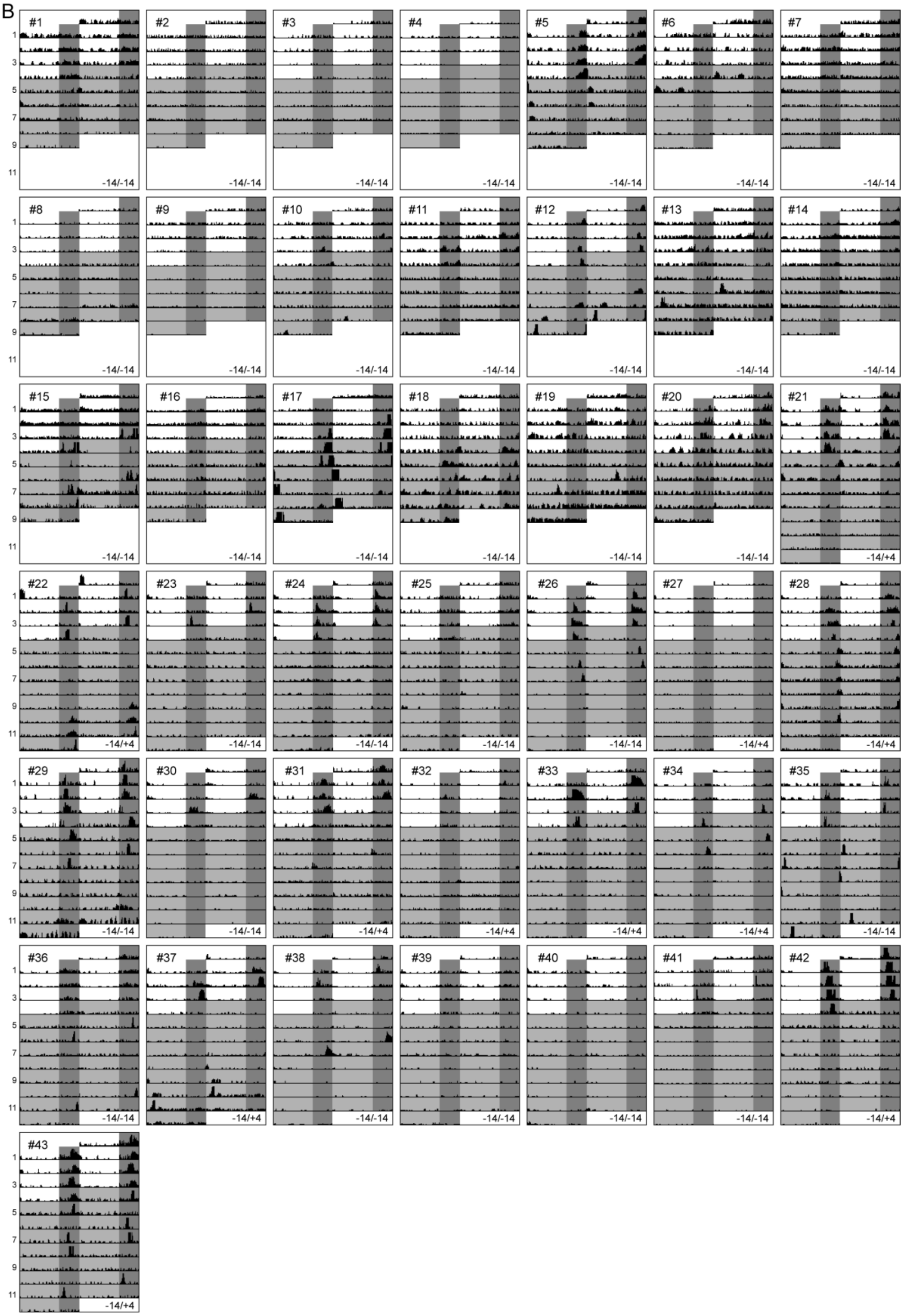

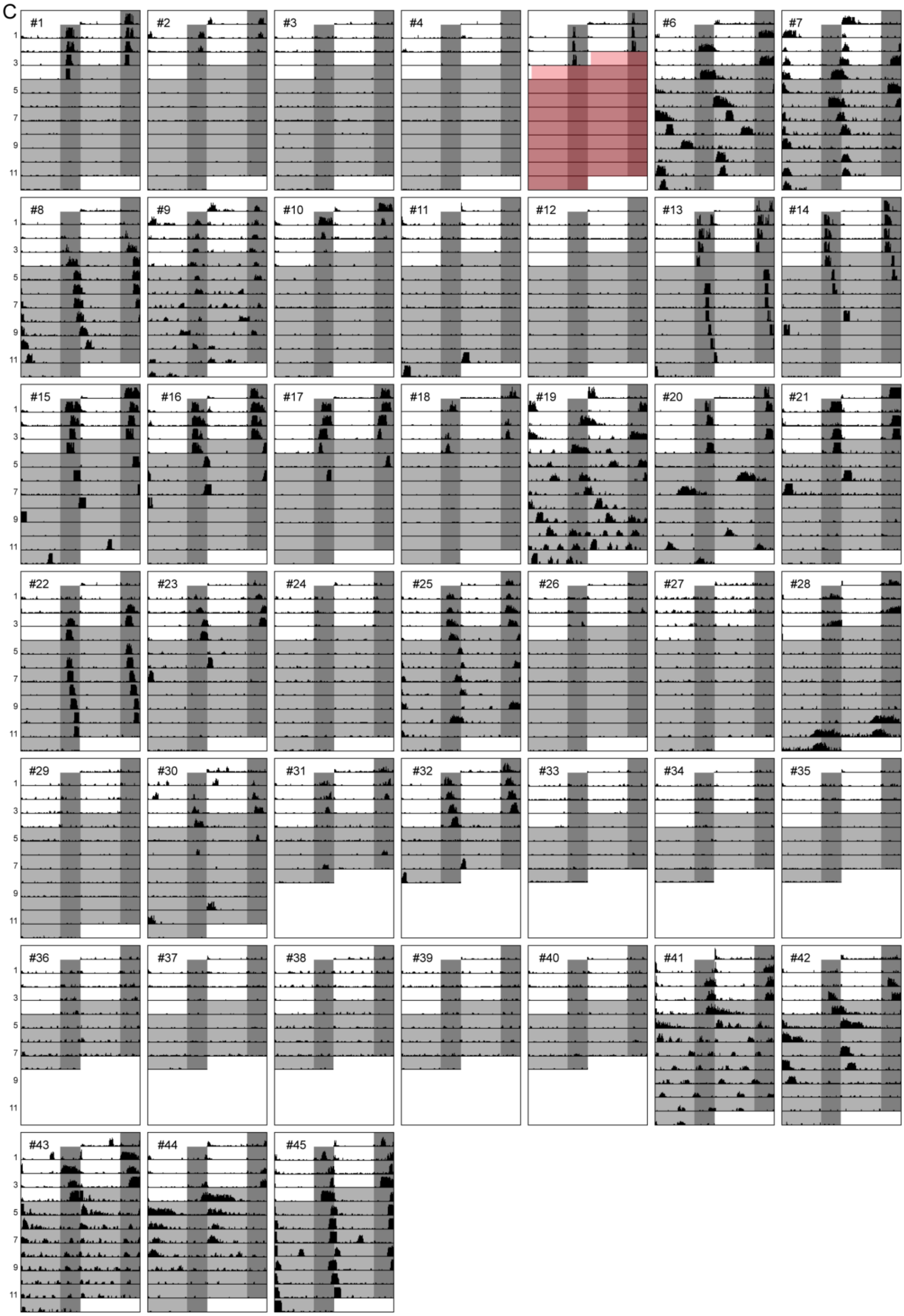
C
#1
#2
#3
#4
#6
#7
#8
#9
#10
#11
#12
#13
#14
#15
#16
#17
#18
#19
#20
#21
#22
#23
#24
#25
#26
#27
#28
#29
#30
#31
#32
#33
#34
#35
#36
#37
#38
#39
#40
#41
#42
#43
#44
#45
1
3
5
7
9
11

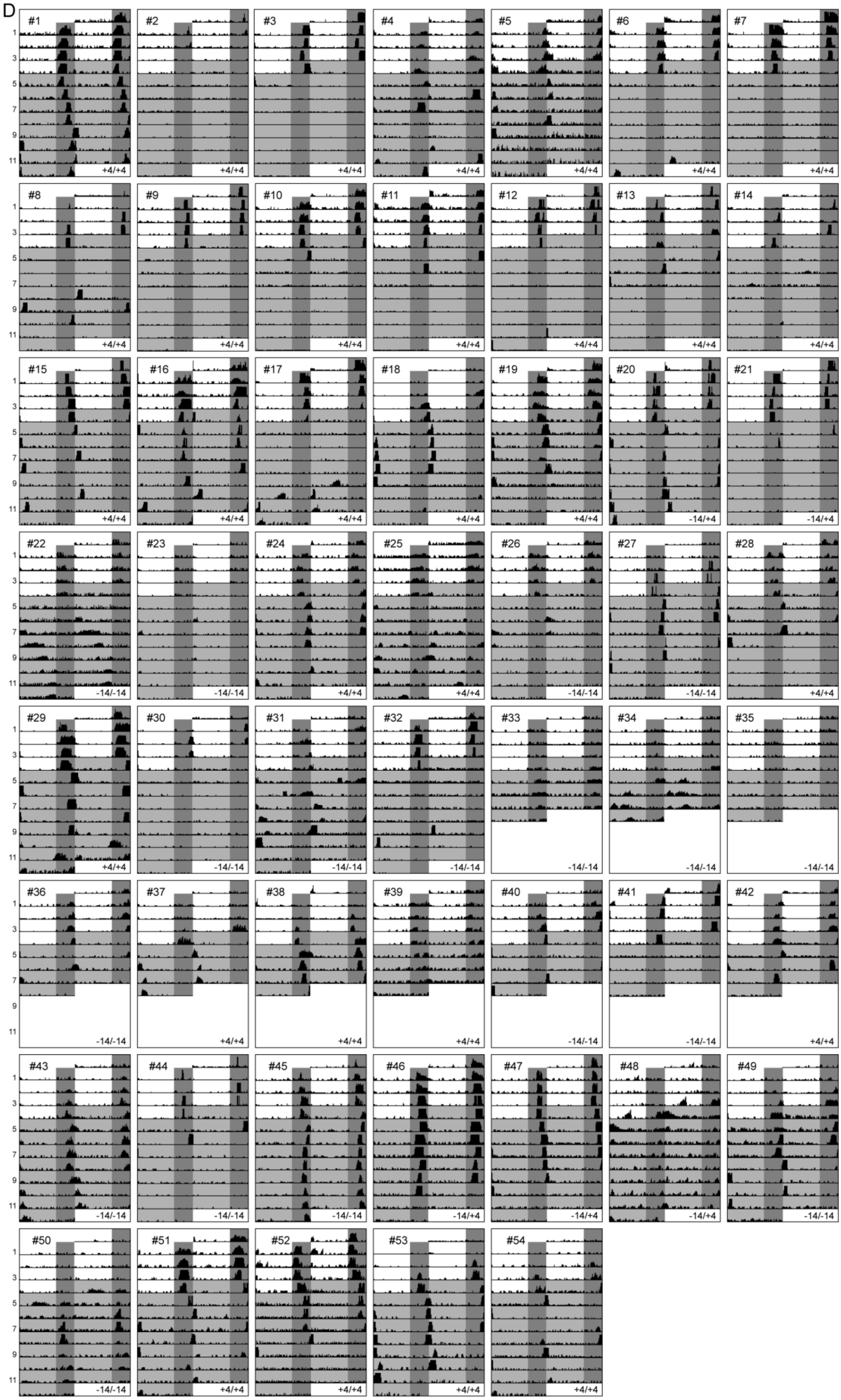
D
#1
#2
#3
#4
#5
#6
#7
#8
#9
#10
#11
#12
#13
#14
#15
#16
#17
#18
#19
#20
#21
#22
#23
#24
#25
#26
#27
#28
#29
#30
#31
#32
#33
#34
#35
#36
#37
#38
#39
#40
#41
#42
#43
#44
#45
#46
#47
#48
#49
#50
#51
#52
#53
#54
+4/+4
-14/+4
-14/-14

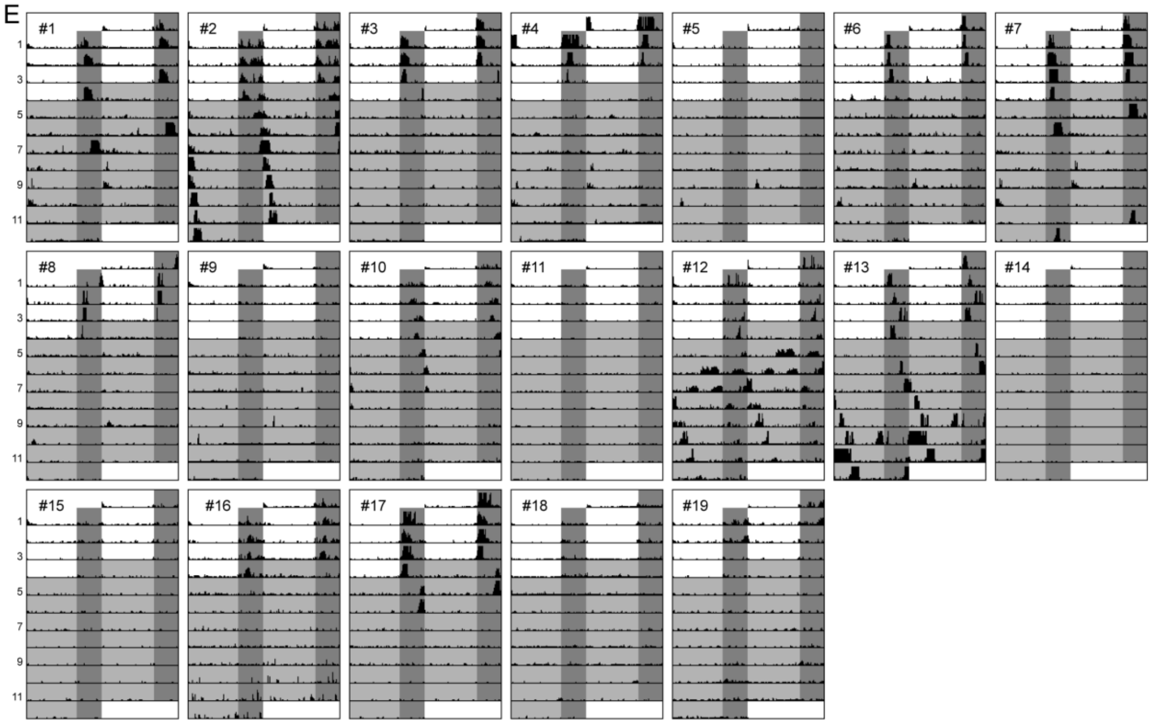
E
#1
#2
#3
#4
#5
#6
#7
#8
#9
#10
#11
#12
#13
#14
#15
#16
#17
#18
#19

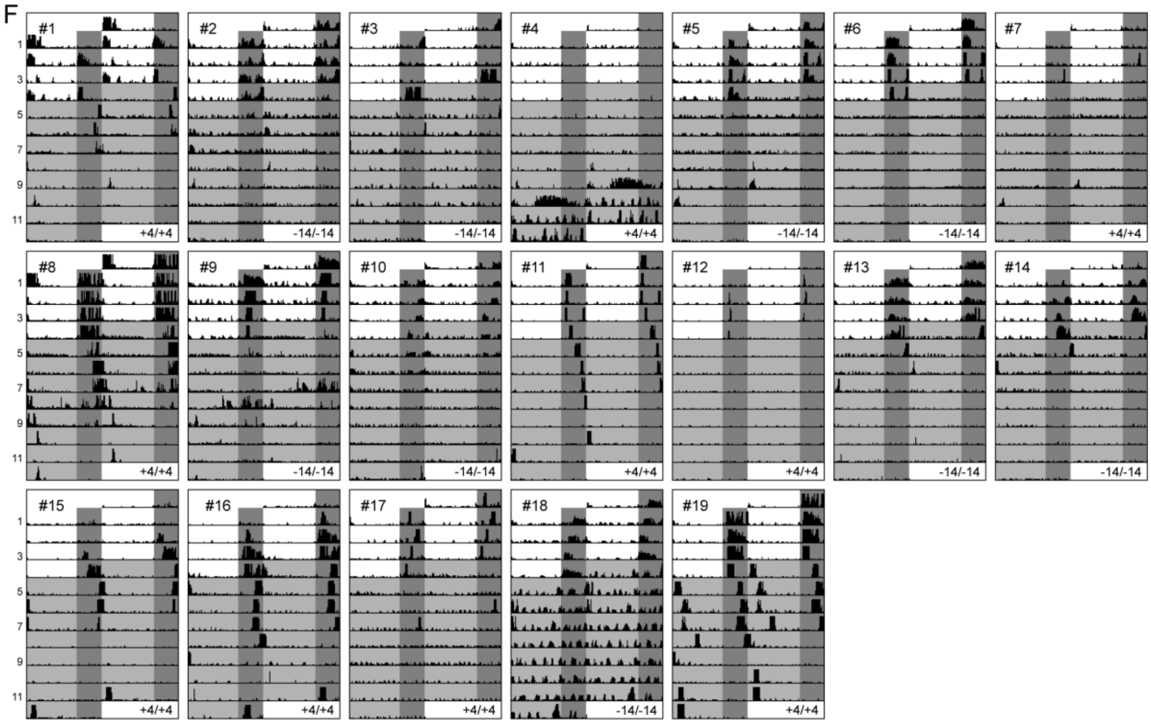
F
#1
+4/+4
#2
-14/-14
#3
-14/-14
#4
+4/+4
#5
-14/-14
#6
-14/-14
#7
+4/+4
#8
+4/+4
#9
-14/-14
#10
-14/-14
#11
+4/+4
#12
+4/+4
#13
-14/-14
#14
-14/-14
#15
+4/+4
#16
+4/+4
#17
+4/+4
#18
-14/-14
#19
+4/+4
